# A dopaminergic visuomotor gateway to the zebrafish optic tectum

**DOI:** 10.64898/2026.09.15.751684

**Authors:** Nils Brehm, Shagnik Chakraborty, Johann H. Bollmann

## Abstract

Dopamine (DA) is widely known as a neuromodulator essential for reward, motivation and learning, yet it also modifies rapid sensorimotor transformations. Understanding how dopaminergic neurons process sensory input and motor-related signals is therefore critical for elucidating their role in sensorimotor control. In vertebrates, a conserved visual center under dopaminergic influence is the superior colliculus, or optic tectum in fish. Here, using anatomical and molecular analyses in larval zebrafish, we identify a defined cluster of pretectal DA (PrDA) neurons whose axons densely innervate the tectum, predominantly in its deep neuropil. Combining functional Ca2+ imaging with visual stimulation and motor recordings, we show that PrDA neurons respond reliably to visual stimulation. However, most PrDA neurons exhibit pronounced activity also during spontaneous locomotion, and enhanced activity when visual stimuli and motor output co-occur, indicating that PrDA neurons integrate convergent input from visual and motor centers. Furthermore, spontaneous PrDA neuron activity was synchronized and the synchrony was even stronger when visual or motor activity contributed to their activation. Notably, as visual stimuli differed in their efficacy to drive swim activity, motor-associated responsiveness of PrDA neurons produced apparent direction selectivity to stimulus motion when motor activity was not accounted for. This apparent neural response bias disappeared once motor activity was taken into account. Together, these findings suggest that PrDA neurons provide rapid, visuomotor-related dopaminergic modulation of tectal circuits that transform visual information into context-dependent motor commands, a principle that may extend to homologous mammalian midbrain circuits

## INTRODUCTION

Dopamine is a conserved neuromodulator with far-ranging influence on neural circuit function across the vertebrate brain. Distinct populations of dopaminergic neurons, organized in anatomically and functionally separable pathways, regulate motor control, reward-based learning, motivation and cognitive flexibility (Bjorklund and Dunnett 2007, Schultz 2007, Klanker et al. 2013, Jin and Costa 2015, Berke 2018, Watabe-Uchida and Uchida 2018). In addition to their roles in reward and movement, several lines of evidence suggest that dopaminergic neurons modulate sensory processing directly, adjusting neuronal excitability and efficacy of synaptic transmission in ascending sensory pathways (Jacob and Nienborg 2018). For example in the auditory pathway, DA acting via D2-like receptors in the subcortical inferior colliculus can selectively suppress neuronal responses to unexpected sounds (Valdes-Baizabal et al. 2020). Likewise, work in the mammalian visual system has shown that dopaminergic inputs in the superior colliculus (SC) can alter neural excitability of SC neurons (Bolton et al. 2015) and modulate the probability of aversive responses to threatening visual stimuli (Montardy et al. 2022). Notably, dopaminergic regulation of visuomotor processing is conserved across vertebrate phyla, as for example in lamprey, dopaminergic neurons project directly to the optic tectum (homologous to the mammalian SC), and modulate neural excitability and behavioral output (Perez-Fernandez et al. 2017). Together, it is clear that dopaminergic signals can strongly influence visuomotor processing in subcortical visual areas such as the superior colliculus/optic tectum (de Malmazet and Tripodi 2023, Ryczko and Dubuc 2023). Yet, it remains unresolved under which stimulus conditions and behavioral states dopaminergic inputs are engaged to adjust visuomotor computations.

The larval zebrafish (*Danio rerio*) has emerged as a useful vertebrate model for studying neural connectivity and circuit function that underlie visually guided behaviors and swift behavioral decisions, owing to its experimental accessibility and compact yet vertebrate-typical brain architecture (Bollmann 2019, Basso et al. 2021, Isa et al. 2021). Moreover, this model offers unique opportunities for dissecting how DA shapes fast visuomotor processing in key visual centers, including the optic tectum. Distinct dopaminergic clusters are located in basal diencephalic nuclei of the posterior tuberculum, with additional dopaminergic populations distributed across the preoptic area, ventral thalamus, pretectum, subpallium, and olfactory bulb (Rink and Wullimann 2002, McLean and Fetcho 2004, Filippi et al. 2010) (Fig. 1A). Recent work has placed these cell groups into an anatomical framework, highlighting conserved organizational principles across vertebrates (Wullimann et al. 2024). Functionally, distinct dopaminergic populations in the hypothalamus and posterior tuberculum have been implicated in rapidly modulating sensory gain and biasing motor circuit excitability in locomotor control (Jay et al. 2015, Yao et al. 2016, Reinig et al. 2017, Barrios et al. 2020, Jha and Thirumalai 2020, Odstrcil et al. 2022). Thus, dopaminergic neurons in zebrafish are likely to provide a set of parallel, partially specialized channels that can influence different steps of sensorimotor transformations, from early sensory encoding to premotor command generation.

**Figure 1:**
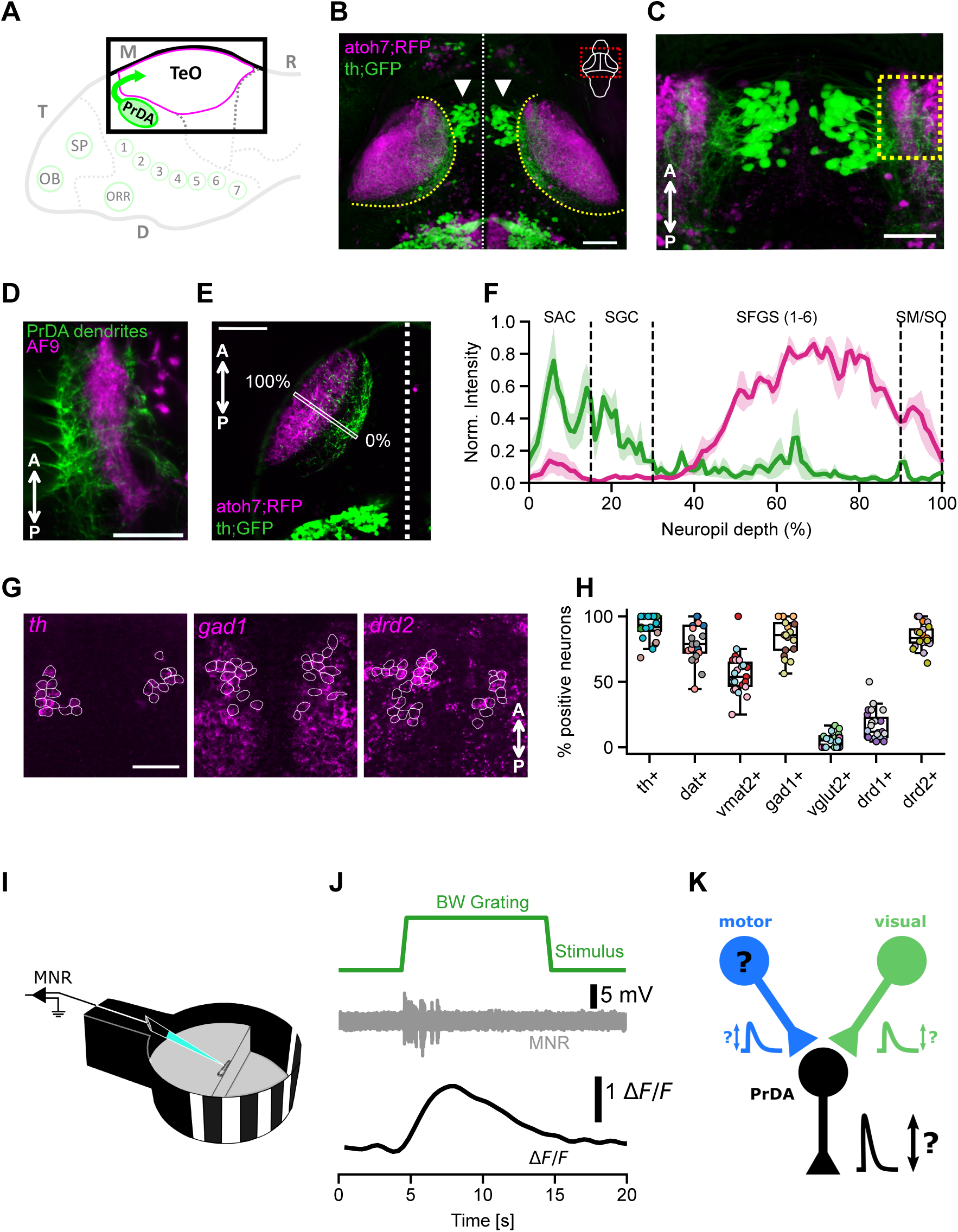
Anatomical organization and *in situ* HCR of marker genes in PrDA neurons. A) Schematic overview of dopaminergic clusters in the larval zebrafish brain. B) Overview (z-projection) of optic tectum and pretectum, showing PrDA neurons (green, arrowheads) and their axons projecting laterally into the tectal neuropil. Tectal neuropil regions identified by RGC axons (magenta). Dashed curve indicates boundary between neuropil and periventricular cell body layer. Scale bar 50 µm. C) Magnified view of PrDA cluster in (B) with dendrites in close proximity to RGC axons. Yellow box: subregion shown in panel D. Scale bar 20 µm. D) Subregion (yellow box in C) of a single optical slice highlighting PrDA dendrites (green) overlapping with RGC axons in AF9 (magenta). Scale bar 20 µm. E) Optical slice of one tectal hemisphere. Rectangle indicates region for measuring intensity profiles of GFP and RFP fluorescence, respectively, across tectal neuropil layers. Scale bar 50 µm. F) GFP and RFP intensity profiles averaged across three larvae. Plots show normalized intensities, with shaded areas showing SEM. G) *In situ* HCR for three marker genes (*th*, *gad1*, *drd2*). Single optical slices show stained gene transcripts (magenta), with ROIs representing somata of *th*-expressing GFP-positive neurons (white outlines). Scale bar 20 µm. H) Fraction of GFP-positive neurons showing HCR-staining for *th*, *dat*, *vmat2*, *gad1*, *vglut2*, *drd1* and *drd2*, respectively (n = 3 larva per marker gene, color coded for specific larvae). I) Experimental setup for simultaneous Ca2+ imaging of PrDA neurons and recording of motor nerve activity. Visual stimuli were presented on a cylindrical screen positioned to the left eye of the larva. J) Example recording of GCaMP6s fluorescence (ΔF/F) from single PrDA neuron during visual stimulation, concurrent with motor nerve activity (MNR, grey trace). Green trace indicates visual stimulation interval with a backward grating. K) Motor activity and visual stimulation as potential drives for PrDA neuron responses. Abbreviations: AF: arborization field, RGC: retinal ganglion cell, SPV: *stratum periventriculare*, SAC: *stratum album centrale*, SGC: *stratum griseum centrale*, SFGS: *stratum fibrosum griseum superficiale*, SO: *stratum opticum*, SM: *stratum marginale*. OB: olfactory bulb, SP: subpallium, ORR: optic recess region, M: mesencephalon, R: rhombencephalon, T: telencephalon, TeO: optic tectum. Numbers 1: ventral thalamic, 2 to 4: rostral and caudal posterior tubercular, 3: medial hypothalamic, 5 and 6: hypothalamic, 7: caudal hypothalamic DA clusters (Kastenhuber et al. 2010, Altbürger et al. 2023).

With respect to dopaminergic innervation of the optic tectum, a comprehensive mapping of dopaminergic projections showed that it arises predominantly from a cluster of neurons in the diencephalic pretectum (Fig. 1A) (Tay et al. 2011, see also Shang et al. 2015). Pretectal cell populations of larval zebrafish are mainly implicated as a hub for optic flow computation, driving stabilizing responses such as the optokinetic and optomotor reflexes, similar to the accessory optic system in mammals (Masseck and Hoffmann 2009, Kubo et al. 2014, Matsuda and Kubo 2021). Consistent with this role, many pretectal neurons exhibit pronounced direction selectivity for optic flow stimuli, with distinct cell populations apparently tuned to different directions of whole-field visual motion (Kubo et al. 2014, Naumann et al. 2016, Wang et al. 2019). Earlier work, however, has not distinguished between dopaminergic and non-dopaminergic neurons within these pretectal populations, leaving open the conditions under which dopaminergic neurons are recruited to modulate sensorimotor transformations in the optic tectum. Furthermore, since the probability and vigor of locomotion in larval zebrafish are themselves sensitive to the direction of visual motion, apparent direction selectivity in pretectal neurons (Kubo et al. 2014, Naumann et al. 2016, Wang et al. 2019), as well as in other visuomotor circuits including the tectum itself, (for example Niell and Smith 2005, Gabriel et al. 2012, Hunter et al. 2013, Förster et al. 2020), could potentially carry a bias from motor-coupled activity rather than representing a purely visual response, a possibility that has not systematically been explored.

Here, we address these questions by characterizing this cluster of dopaminergic neurons in the pretectum (‘PrDA neurons’) in a recently established zebrafish line in which dopaminergic neurons throughout the larval brain are robustly labeled (Altbürger et al. 2023). We map their somatic location and long-range projections, revealing specific innervation of deep tectal layers. By monitoring their activity during stimulation and spontaneous locomotion, we show that these PrDA neurons are transiently activated by a broad range of visual stimuli and are additionally recruited during motor output, independent of visual drive. Together, these results suggest that PrDA neurons exert dynamic, stimulus- and locomotor-dependent modulatory control over tectal visuomotor processing, pointing to an evolutionarily conserved motif shared with dopaminergic modulation of the mammalian superior colliculus.

## RESULTS

### Anatomy and molecular identity of PrDA neurons

To characterize the anatomy of pretectal dopaminergic neurons, we used the *tyrosine hydroxylase*-(*th*)-specific QF2-driver line *Tg(th:th-e2A-QF2)m1512* (Altbürger et al. 2023), crossed with the reporter line *Tg(QUAS-GFP)c403* (Subedi et al. 2014), enabling Q-system-dependent labeling of pretectal neurons shown previously to be dopaminergic. To relate their position and projections to optic tectum anatomy, we imaged GFP-positive PrDA neurons in *Tg(th:th-e2A-QF2);Tg(QUAS-GFP);Tg(atoh7:Gal4-VP16)*;*Tg(UAS:RFP)* transgenic larvae, in which additional RFP expression in retinal ganglion cells (RGC) and their axons labels the retinorecipient layers of the tectum and other RGC arborization fields (Fig. 1B-D).

PrDA somata were located medioventral to the anterior pole of the optic tectum and formed two compact clusters flanking the midline, with axons projecting into the tectal neuropil (Fig. 1B,C). The dendrites of PrDA neurons overlapped with RGC axons in arborization field 9 (AF9, Fig. 1D), suggesting potential direct retinal input. Previous work showed that PrDA axons project to the optic tectum (Tay et al. 2011), but their laminar organization relative to the retinorecipient layers remained unclear. In single optical sections of tectal hemispheres we found a stratified pretecto-tectal projection pattern (Fig. 1E). Quantification across neuropil depth showed that PrDA axons were most abundant in the deeper SAC and SGC layers, extended sparsely into the retinorecipient SFGS, and were rarely present in the superficial SM and SO layers (Fig. 1F). Thus, PrDA projections predominantly target deeper tectal layers while partially overlapping with retinal inputs in the SFGS.

We next characterized PrDA neurons molecularly using *in situ* hybridization chain reaction (HCR; Fig. 1G,H and Suppl. Fig. 1). Most GFP-positive neurons expressed the dopaminergic marker genes *th*, *dat*, and *vmat2*, confirming their dopaminergic identity. Given that dopaminergic neurons can co-release classical neurotransmitters (Granger et al. 2017), we assessed markers for glutamatergic and GABAergic transmission. The majority of PrDA neurons were *gad1*-positive (median: 85%; n = 3 larvae; Fig. 1G,H), consistent with previous work (Filippi et al. 2014), whereas *vglut2* expression was negligible (median: 5%; n = 3 larvae; Fig. 1H). Finally, we examined dopamine receptor expression and found that few PrDA neurons expressed *drd1*, while a large fraction expressed *drd2* (median: 11% and 83%, respectively; n = 3 larvae; Fig. 1G,H), consistent with a potential role for D2-type-autoreceptor-mediated feedback that may regulate their activity and transmitter release.

Taken together, PrDA neurons are clearly positioned to modulate tectal visuomotor processing, however the conditions under which these neurons are active are not well understood.

### Sensory and motor contributions to PrDA neuron activity

Therefore, to address their functional role, we imaged PrDA neuron activity in *Tg(th:th-e2A-QF2); Tg(QUAS:GCaMP6s)* transgenic larvae, in which GCaMP6s is robustly expressed in these dopaminergic neurons, while presenting visual stimuli and concurrently recording fictive motor output (Fig.1I). PrDA neurons exhibited pronounced activity when, for example, a moving grating was presented, suggesting that these neurons could be driven by visual stimulation (Fig. 1J). However, it is difficult to distinguish whether this response is directly caused by visually evoked synaptic input or rather by motor-associated excitatory drive (Fig. 1K), as many visual stimuli also elicit locomotion. To discriminate between these possibilities and to classify their functional properties, we recorded PrDA neuronal activity and fictive swimming behavior simultaneously while presenting a battery of visual stimuli, including moving targets, gratings, looms, and flashes (Fig. 2A-C). During Ca2+ imaging, immobilized larvae exhibited robust fictive swimming, evident as burst activity in the motor nerve recording (MNR, Fig. 2C, grey trace), which we used to detect the onset and offset of fictive swim events (Suppl. Fig. 2A) (Masino and Fetcho 2005, Ahrens et al. 2012). Then, to quantify neuronal responses of all recorded neurons (Suppl. Fig. 2B) and to relate them to visual stimuli and motor events, we used a linear-regression-based response scoring approach, using an idealized Ca2+ response trace (Suppl. Fig. 2C-F) and the intervals of visual stimulation and motor activity to construct stimulus and motor regressors, respectively (Fig. 2C). By comparing these to the measured fluorescence signals for each PrDA neuron, we obtained response scores for each visual stimulus presentation as well as for each detected swim event (Fig. 2C, bottom traces, and Suppl. Fig. 2G-I).

**Figure 2:**
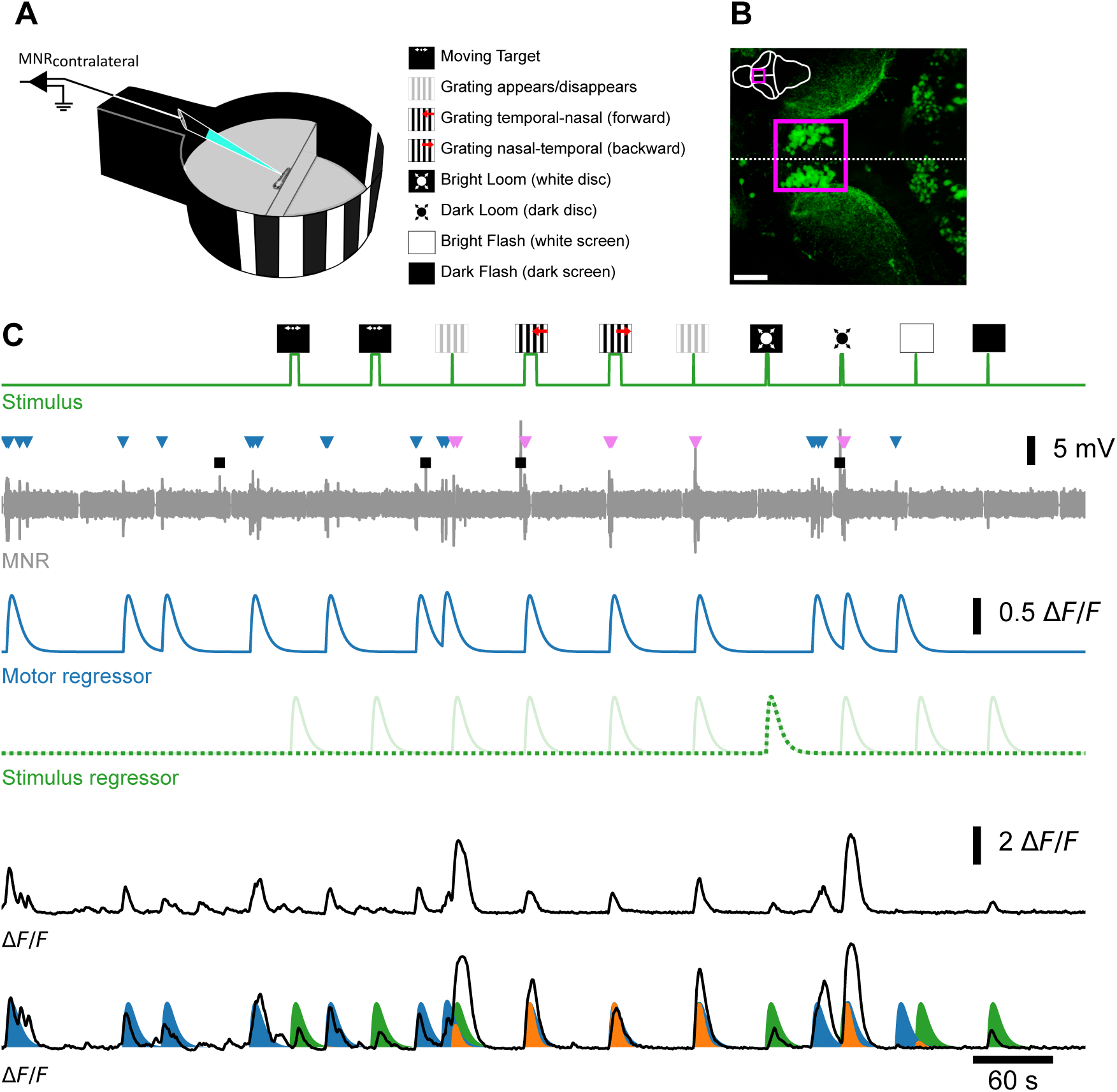
Functional imaging of PrDA neurons. A) Experimental setup and description of different visual stimulus types. B) Overview (Z-projection) of optic tectum and pretectum. Magenta rectangle depicts the scan area for recording GCaMP6s fluorescence time course in PrDA neurons. Scale bar 50 µm. C) Example recording. From top to bottom: stimulus trace (green) with pictograms indicating stimulus type; MNR trace (gray) with spontaneous swim bouts (blue arrowheads), stimulus-associated swim bouts (magenta arrowheads) and recording artifacts not corresponding to swim bouts (black rectangles). The motor (blue) and stimulus (green) regressors used for response quantification are shown below. For visualization, the regressor corresponding to the bright loom stimulus is highlighted with a dotted line. Two example ΔF/F traces from different neurons are shown in black. In the second trace, motor (blue) and stimulus (green) regressors are overlaid; coincident events are shown in orange.

### Fictive locomotor output strongly correlates with PrDA neuron activity

First, to isolate the contribution of motor activity, we focused on spontaneous fictive swim bouts and excluded all motor events that occurred in close temporal proximity to visual stimuli. For each neuron, we computed response scores to the remaining spontaneous swim events and assessed their significance with a bootstrap procedure (Suppl. Fig. 2J,K), classifying neurons as responsive or non-responsive (representative neuron shown in Fig. 3A). For robustness, we restricted this analysis to sweeps containing at least three spontaneous swim events. Swim-associated activity was relatively reliable across the PrDA population: in 75% of recorded neurons, at least 37% of spontaneous swim events were accompanied by a significant Ca2+ response and half of all neurons had a response probability of 50% or more (Fig. 3B). Motor-related responsiveness was broadly distributed across the PrDA population, with no apparent spatial clustering (Fig. 3C), pointing to a functionally homogeneous representation of motor input. Together, these findings show that PrDA neurons are strongly and reliably activated during spontaneous motor events.

**Figure 3:**
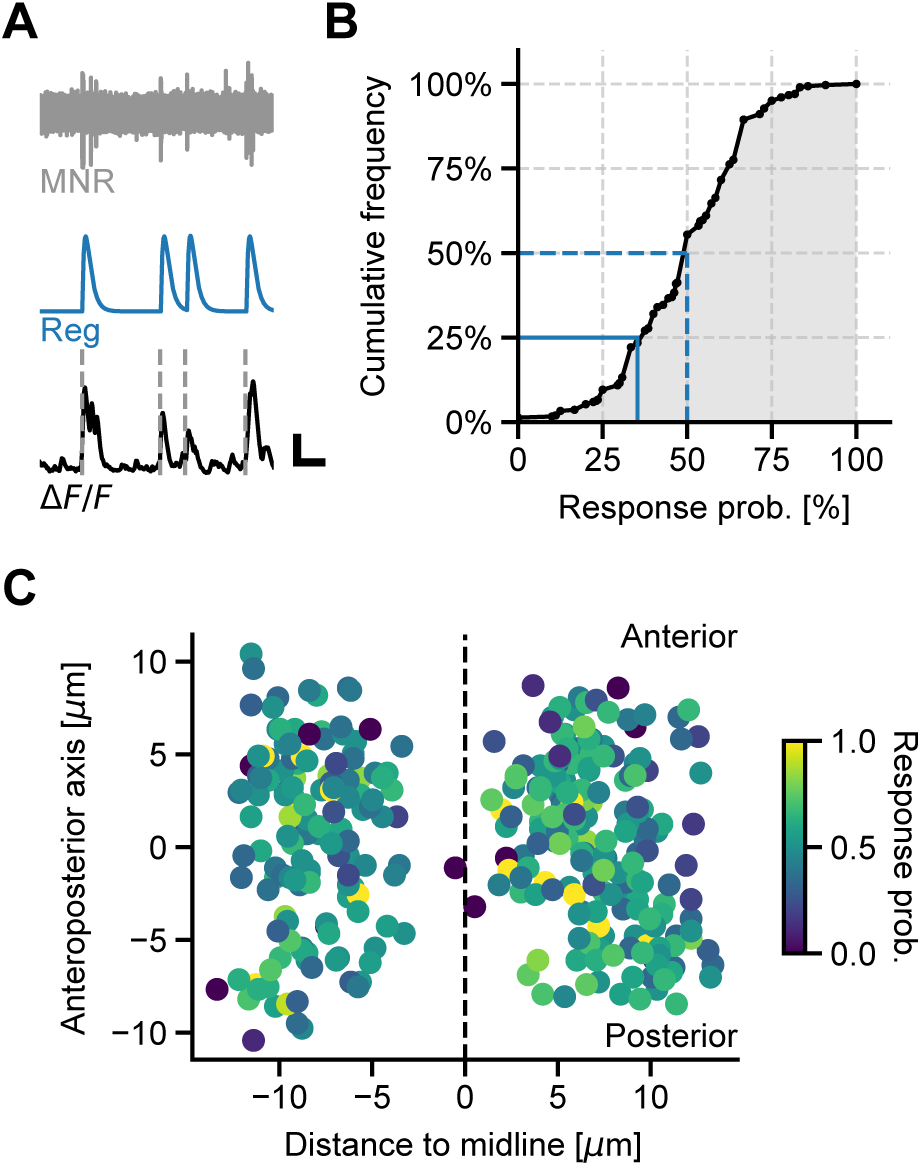
Spontaneous motor events are associated with phasic PrDA neuron activity. A) Example recording showing MNR trace (top), motor regressor trace (middle), and ΔF/F trace (bottom). Dashed vertical lines indicate significant neural responses. Scale bars: 1 ΔF/F and 30 seconds. B) Cumulative distribution of the response probability associated with spontaneous motor events per neuron and sweep (n = 303 neurons). The ratio between the number of significant responses and the total number of spontaneous motor events defines the response probability of a neuron. Only sweeps with at least 3 spontaneous motor events were included. Blue solid and dashed lines mark 25% and 50% frequency value, respectively. C) Spatial distribution of all neurons with response probability color-coded. Same data as in B (n = 303).

### PrDA neurons are driven by visual stimulation and respond bilaterally

We next examined how PrDA neurons respond to visual stimulation. Different visual stimuli varied markedly in how reliably they evoked swimming. Dark looms and forward-moving gratings were the strongest drivers of motor activity, whereas bright flashes and bright looms elicited swims only occasionally (Suppl. Fig. 3). At the same time, many PrDA neurons were phasically active also during visual stimulation alone, that is, when the stimulus did not evoke swimming, raising the possibility that motor and visual contributions are entangled in these neurons.

Comparing the fluorescence traces from all recorded neurons (Fig. 4A; for responses from individual neurons see Fig. 4C), while marking concurrent motor events (blue bars, Fig. 4A, bottom), we found that motor activity alone did not suffice to explain PrDA neuron activity during visual stimulation. To demonstrate this, we attenuated the fluorescence signals using the motor regressors (Fig.4B), generating a motor-corrected view of the data. After strongly suppressing the motor component, purely visual responses became more apparent, confirming that PrDA neurons can indeed be visually driven independent of motor activity (Fig. 4B, bottom). On the other hand, when comparing the responses of PrDA neurons to, for example, bright and dark looms, we found that ignoring the motor component inflated the apparent visual response amplitude and did so more for dark looms, since this stimulus is more likely to elicit a swim event (Fig. 4D and Suppl. Fig. 3).

**Figure 4:**
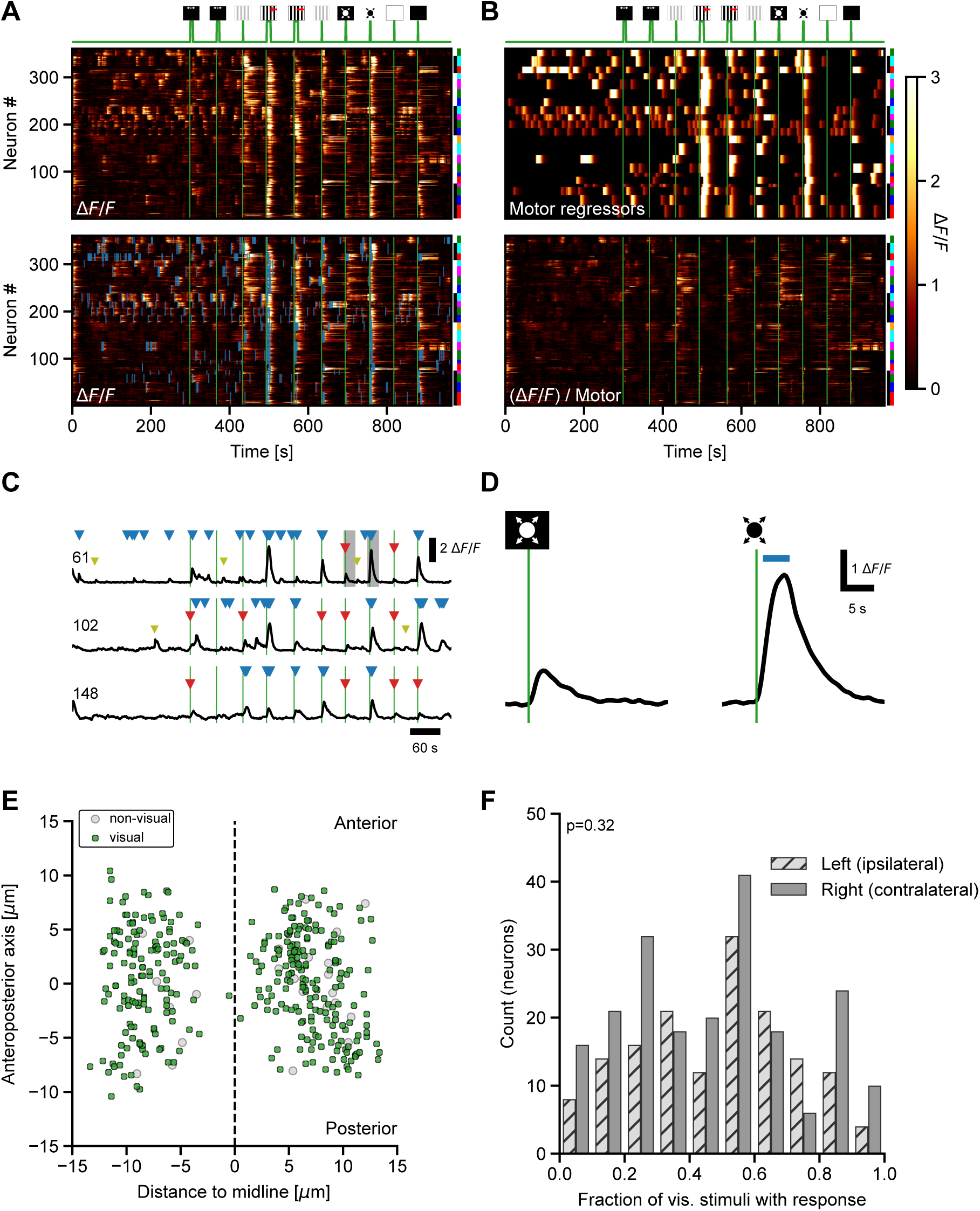
PrDA neurons are driven by visual stimulation and respond bilaterally. A) Activity heat map of PrDA neurons (same color scale as in B). Visual stimulation is indicated by the green trace and pictograms above. Bottom: Same heat map, but with motor events indicated by blue bars (n = 360 neurons, 6 larvae). Larva ID is indicated with black and white stripes and z-plane is color coded. B) Activity heat maps of motor regressors (ΔF/F color-coded) for the neurons and sweeps in A). Bottom: Measured activity heat map from A, attenuated by motor regressors, emphasizing the contribution of visual stimulation to the overall responses of PrDA neurons. C) Example ΔF/F traces (neuron IDs correspond to those shown in A). Green lines mark visual stimulus onsets (as in A and B). Olive arrows show spontaneous activity, blue arrows indicate swim bouts, and red arrows indicate purely visual responses (stimulus without accompanying swim). D) Zoomed-in traces from C (gray rectangles in top trace) for the ΔF/F response to a bright loom (left) and a dark loom (right). Blue bar indicates motor activity. E) Spatial distribution of all neurons with either no significant visual response (gray) or with at least one significant visual response (green) in the absence of motor events. F) Histograms of the fraction of stimulus types that resulted in a significant response in the absence of motor events for all neurons, separated by left and right pretectal hemisphere, ipsi- and contralateral to the stimulated eye, respectively. No difference between left and right hemispheres (Mann–Whitney U test, U = 16822, p = 0.32, n=6 fish).

Mapping the spatial distribution of PrDA neurons that exhibited significant activity in response to visual stimulation in the absence of swimming, we found no anatomical subregion enriched for such neurons (Fig. 4E). Since we stimulated only the left eye and RGC axons project exclusively to the contralateral hemisphere, the left (ipsilateral) hemisphere received no direct retinal input. Nonetheless, the majority of ipsilateral PrDA neurons responded significantly to multiple visual stimulus types, with no difference in the width of responsiveness between the left and the right hemisphere (Mann–Whitney U test, U = 16822, p = 0.32, n=6 fish; Fig. 4F). Therefore, monocular visual information is represented bilaterally in PrDA neurons, and direct RGC input cannot be the sole source for visually driven responses in PrDA neurons.

### PrDA neurons combine motor and visual inputs

To assess how PrDA neurons integrate the combined influence of visual stimulation and concurrent motor activity, we quantified responses for each neuron using both motor and visual regressors (prefixes ‘m’ and ‘v’, respectively). As noted above, stimulus types differed strongly in how reliably they triggered a swim response. For dark looms, which almost always triggered a swim, the visual and motor components were inseparable, as reflected in the close similarity between stimulus and motor scores across neurons (mDL and vDL, Fig. 5A). In contrast, bright looms triggered fewer swims, but many neurons were still active in the absence of a motor event (mBL and vBL, Fig. 5A). Indeed, 158 of 360 neurons showed significant visual response scores even though the larva did not swim during the bright loom. A similar pattern occurred for gratings, where forward-moving gratings reliably drove swimming and therefore produced nearly identical visual and motor scores (mGF and vGF, Fig. 5A). More generally, for stimulus types that did not reliably evoke fictive motor output (e.g. bright flash, BF; bright loom, BL; grating appears, GA), significant visual responses were observed in many neurons, confirming that PrDA neurons can be driven by visual stimulation alone.

**Figure 5:**
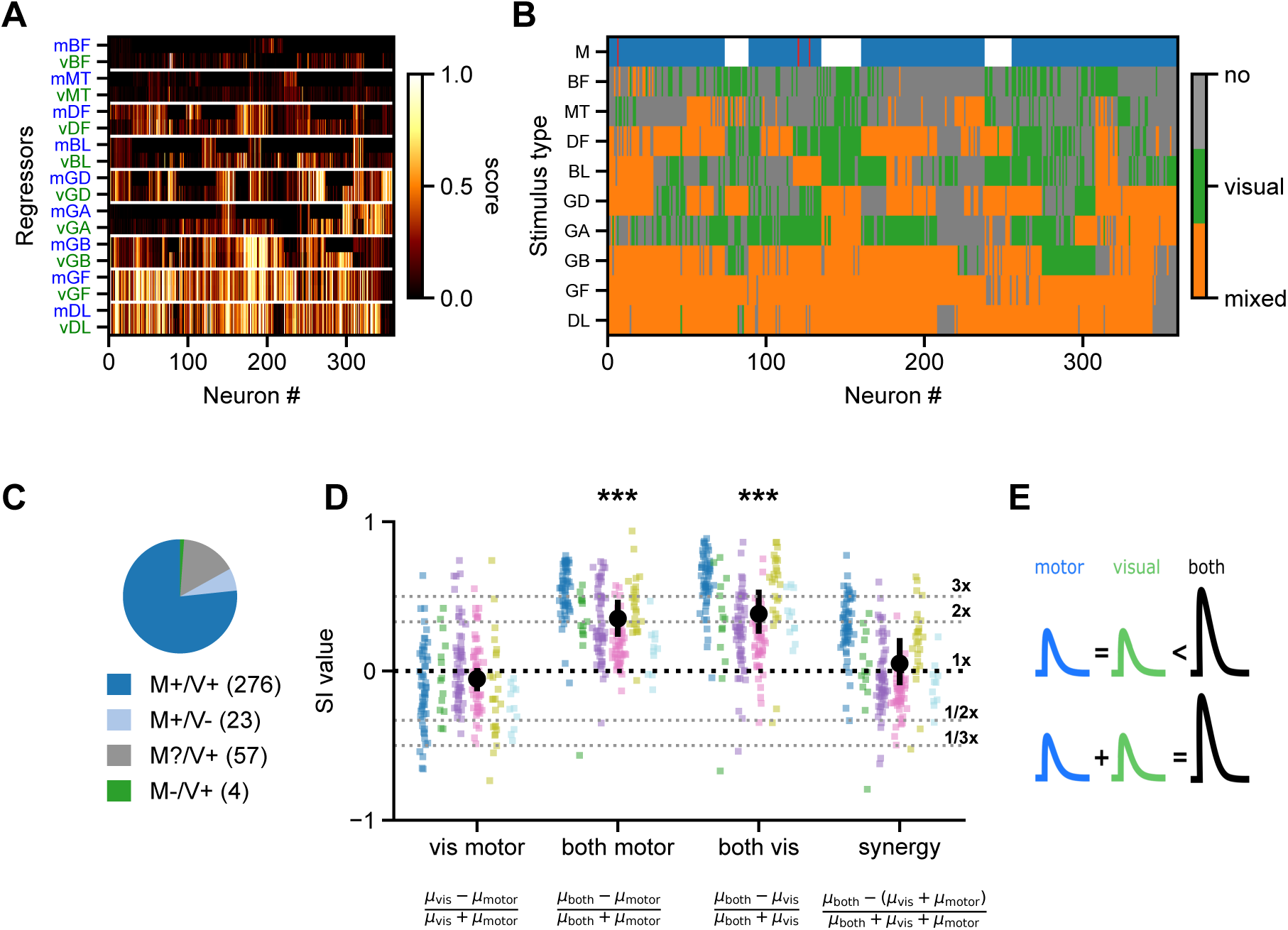
PrDA neurons combine motor and visual inputs. A) Normalized stimulus regressor scores and motor regressor scores for all stimulus types and neurons, sorted from lowest to highest overall score value averaged across neurons (top to bottom). Color scale represents normalized score magnitude. Row labels show stimulus types, with motor (m) and visual (v) regressors marked in blue and green respectively. B) Classification matrix summarizing response scores for all stimulus types. Neurons with a significant score to a stimulus type in the absence of a motor event are depicted in green (i.e. purely visual), otherwise in orange (mixed). Neurons with no significant responses are shown in gray (no response). The top row labeled “M” indicates responsiveness to spontaneous motor events: blue denotes neurons with activity associated with spontaneous motor events; red indicates neurons without significant responses during spontaneous motor events, but during visual stimulation; and white denotes neurons with insufficient data (sweeps with <3 spontaneous motor events). C) Proportion of PrDA neurons classified by their motor and visual drive, respectively. M+: motor-driven; M-: not motor-driven; M?: not determined; V+: visually driven; V-: not visually driven. The number of neurons in each class is shown in parentheses. D) Signed selectivity indices (SI). Colored dots represent individual neurons grouped by larva. Black dots indicate the across-larva mean, with error bars showing the confidence interval. Black dotted line marks zero, corresponding to no difference. Grey dotted lines indicate the ratios of the first mean µ over the second µ for each SI, respectively (3x, 2x, 1/2x. 1/3x). The corresponding equations to calculate SIs are stated below for each condition. E) Schematic drawing highlighting the results shown in D. Top row: across the population, PrDA neurons respond equally to motor activity and visual stimuli (blue and green, respectively), and their activity is enhanced when motor and visual drive coincide (‘both’). Bottom row: Overall, activity in PrDA neurons during motor events in the presence of visual stimulation is the linear sum of the separate, motor-associated and visually driven components, respectively. Stimulus abbreviations: DL, Dark Loom; GF, Grating Forward; GB, Grating Backward; GA, Grating Appears; GD, Grating Disappears; BL, Bright Loom; DF, Dark Flash; MT, Moving Target; BF, Bright Flash; M, Responsiveness to spontaneous motor events.

Next, to visualize the distribution of responses to motor events and visual stimulation across the recorded population, we classified neurons using an index that compared stimulus-driven and motor-associated response scores for every stimulus type (Fig. 5B; neurons were labeled ‘no’, ‘visual’, or ‘mixed’ response, grey/green/orange, respectively). At the level of spontaneous motor activity (top row, M, Fig. 5B), only four neurons were inactive during spontaneous motor events (red bars, M-/V+; Fig. 5B,C). All remaining neurons were either significantly motor-associated (blue bars) or lacked sufficient data for classification (that is, recorded in a sweep with fewer than 3 spontaneous swims, white bars in Fig. 5B). Across all stimulus types—except dark loom and forward grating, where visual and motor signals were inseparable—neurons with significant visual responses were present.

Because the majority (>75%) of neurons exhibited activity both during visual stimulation and spontaneous motor events (Fig. 5C), we next asked how these two putative input signals are summed when they occur together. To analyze this integration, we computed signed selectivity indices from the mean scores of three conditions: visual stimulation without concurrent motor activity (‘visual’), spontaneous motor events (‘motor’) and visual stimulation accompanied by motor activity (‘both’). Neurons for which not all three conditions were met were excluded (57 of 360, M?/V+, Fig. 5C). Each index ranges from −1 to +1 and quantifies the relative preference between two conditions, with positive values indicating a stronger response for the first. The visual-motor index was centered near zero across animals, indicating that, at the population level, neurons respond similarly to visual and motor input when each was present alone (‘vis motor’, Fig. 5D). By contrast, the both-vs-motor and the both-vs-visual indices were significantly greater than zero for most neurons (p < 0.0001, hierarchical bootstrap), showing that simultaneous visual and motor input nearly always evoked stronger responses than either of them alone (“both motor” category and “both vis”, Fig. 5D).

Finally, we computed a synergy index to quantify how the co-occurring motor-associated and visually driven signals summed, i.e. whether the combined response was larger or smaller than the sum of the two individual responses. Across animals, the mean synergy index did not differ significantly from zero (p=0.6, hierarchical bootstrap), indicating that that summation was approximately linear on average (“synergy”, Fig. 5D). However, synergy indices for individual cells ranged from sub-linear to supra-linear, raising the possibility that PrDA neurons integrate visual and motor inputs in diverse ways and could represent a balanced mixture of computations.

In summary, PrDA neurons responded comparably to motor and visual input presented alone, while their activity was increased when the two coincided. At the population level, this elevated activity reflected the linear sum of the individual inputs (see schematic summary in Fig. 5E).

### PrDA neurons exhibit strong synchronized activity

Next, to estimate whether PrDA neurons operate as a homogeneous population to signal relevant events (that is, visual stimuli, locomotion) to the optic tectum, we quantified the degree of synchronization among PrDA neurons. We computed pairwise cross-correlation of z-scored activity across all neurons in each recorded sweep. To disentangle the contribution of visually driven and motor-associated input to this synchrony, we computed cross-correlations separately for time windows that included or excluded these events. Across conditions, the average cross-correlation peaked at zero lag, indicating that PrDA neurons co-activated synchronously rather than with a consistent temporal offset (Fig. 6A). Notably, synchrony was already evident during baseline spontaneous activity, in the absence of any events (Baseline, Fig. 6A). The introduction of relevant events further elevated this synchrony. The strongest synchronization occurred when visual stimuli were accompanied by a swim response, although spontaneous swims and visual stimuli alone also raised correlations substantially above baseline (Fig. 6A). Randomly shuffling the temporal alignment reduced correlations to chance level (Shuffled, Fig. 6A), confirming that the observed synchrony reflected genuine temporal structure.

**Figure 6:**
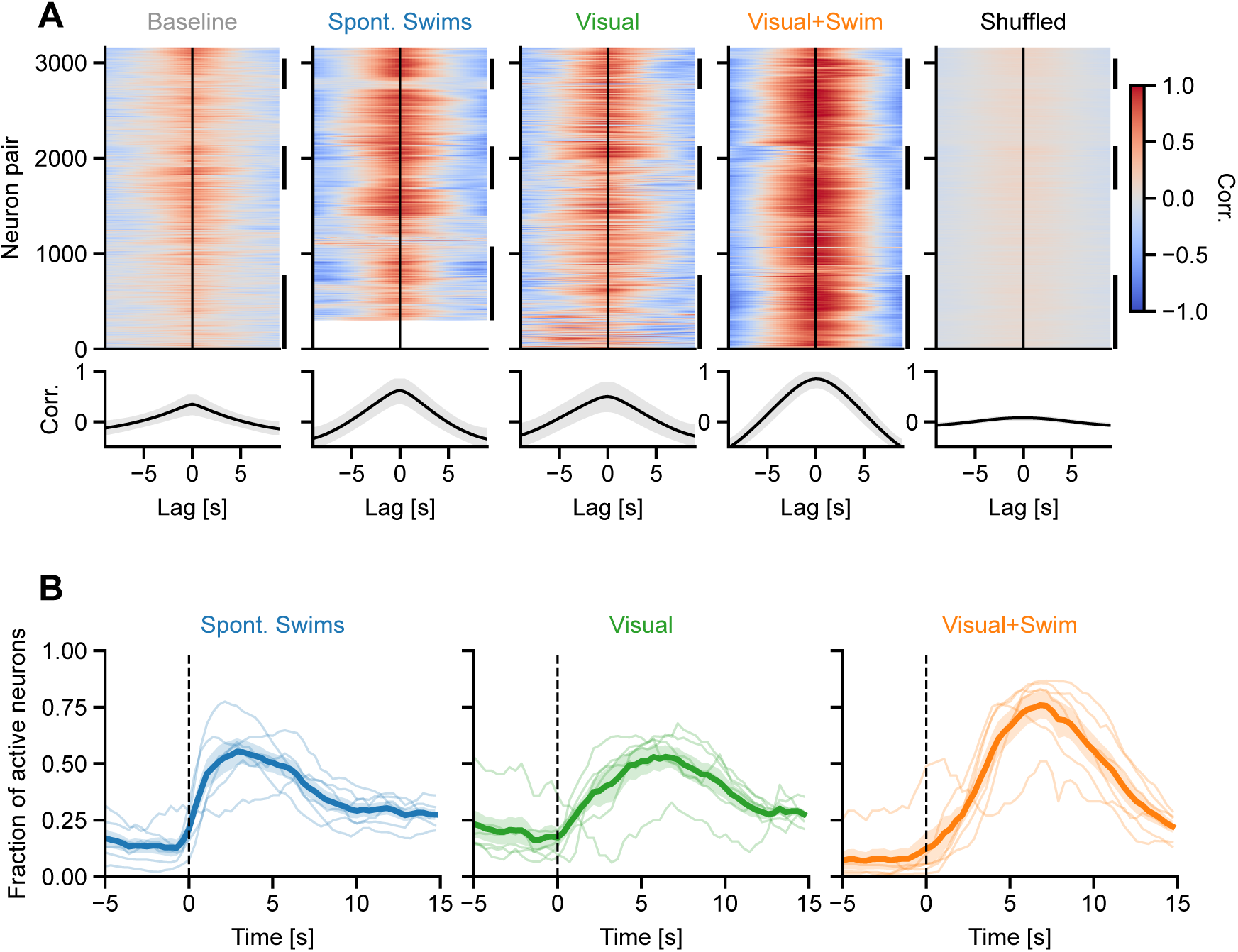
PrDA neurons exhibit strong synchronized activity. A) Cross-correlation heat maps based on the z-scored activity trace of different windowed conditions. The value of the correlation is color coded. Below each heat map, the average correlation over lags is indicated (mean ± standard deviation). The missing neuron pairs for spontaneous swims (2^nd^ panel) is due to one sweep not having any spontaneous swim events at all. B) Recruitment of neurons at the population level as the fraction of active neurons averaged across larvae for spontaneous swims (blue), visual stimuli without a swim response (green) and visual stimuli with a swim response (orange). Thin lines indicate individual larva (n=6); bold lines show mean ± SEM. Vertical dotted line indicates the onset of events.

We next examined the temporal dynamics of the recruitment of neurons at the population level by looking at the fraction of neurons that are active at each time point. Because this measure is aligned to event onset, we quantified it for the three event types rather than for baseline or shuffled conditions (n = 6 larvae, thin lines in Fig. 6B). Following spontaneous swim onset, the fraction of active neurons rose rapidly, peaking at 55% (blue, Fig. 6B). Similarly, during visual stimuli not accompanied by a swim response, the fraction of active neurons peaked at 53% (green, Fig. 6B). When visual stimuli and swim responses co-occurred, the recruitment exceeded either condition alone, reaching 76% of simultaneously active neurons (orange, Fig. 6B). In summary, PrDA neurons were synchronized in their baseline activity and the synchrony was even stronger when there were additional input drives.

### Apparent stimulus selectivity of PrDA neurons is explained by motor activity

Visually responsive neurons are commonly classified according to their stimulus selectivity; that is, the extent to which their responses vary along a particular stimulus dimension, such as stimulus direction or orientation. However, when a neuronal population also responds to motor events, estimates of stimulus selectivity that do not account for motor-related activity may be inaccurate. To assess the extent to which this can bias the interpretation of neuronal response properties, we calculated direction selectivity indices (DSIs) for PrDA neuron responses to forward and backward grating motion and compared the resulting DSI distributions with and without accounting for motor activity.

As expected, the behavioral DSI, based on the number of swims elicited by forward versus backward gratings was shifted towards forward motion (median DSI_swim =_ 0.43; Wilcoxon signed-rank test vs. zero: W = 1781, p < 0.001) and its distribution differed significantly from uniform (KS test: D = 0.35, p < 0.001; Fig. 7A). This reflects the notion that forward gratings triggered more swim responses than backward gratings over all recordings.

**Figure 7:**
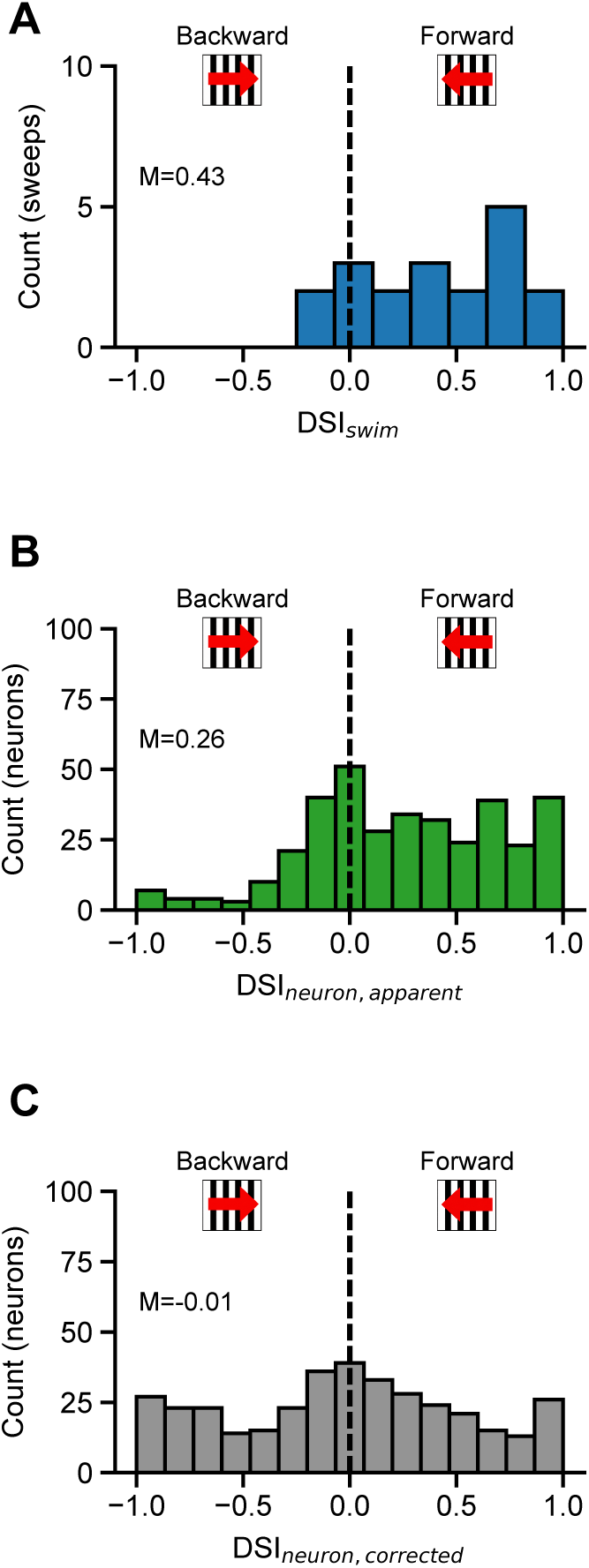
Correcting direction selectivity of PrDA neurons by including motor activity. A) Direction selectivity index distribution based on a larva’s swim response to forward and backward gratings (median DSI_swim =_ 0.43; Wilcoxon signed-rank test vs. zero: W = 1781, p < 0.001; KS uniform test: D = 0.35, p < 0.001; 6 larvae and 21 sweeps) B) Direction selectivity index distribution based on stimulus regressor scores for forward and backward gratings of all neurons (median DSI_neuron,apparent =_ 0.26; Wilcoxon: W = 14508, p < 0.001; KS uniform test: D = 0.27, p < 0.001; 6 larvae, 21 sweeps and 360 neurons). C) Direction selectivity index distribution based on swim-corrected stimulus regressor scores for forward and backward gratings of all neurons (n=360). Each score was attenuated by the number of swim bouts present during visual stimulation (median DSI_neuron,corrected =_ −0.01; Wilcoxon: W = 31747, p = 0.71; KS uniform test: D = 0.07, p = 0.06; 6 larvae, 21 sweeps and 360 neurons).

A qualitatively similar forward shift was present in the apparent neuronal DSI, computed from the stimulus regressor scores for forward and backward gratings (median DSI_neuron,apparent =_ 0.26; Wilcoxon: W = 14508, p < 0.001; KS: D = 0.27, p < 0.001; Fig. 7B). This apparent neural selectivity for forward visual motion mirrors the behavioral bias (Fig. 7A), suggesting that it reflects differences in swim-associated activity rather than a genuine neural preference for forward grating motion.

To test this, we corrected each stimulus score for the number of swim bouts occurring during visual stimulation. The motor-corrected neuronal DSI was no longer shifted forwards, the distribution was centered on zero (median DSI_neuron,corrected =_ −0.01; Wilcoxon: W = 31747, p = 0.71) and it was not significantly different from a uniform distribution anymore (KS: D = 0.07, p = 0.06; Fig. 7C). Thus, the apparent forward direction selectivity of PrDA neurons is largely accounted for by motor activity rather than by tuning of PrDA neuron responses to a particular visual motion direction.

## DISCUSSION

PrDA neurons establish a mixed visuomotor signal that scales in magnitude with combined visual and motor drive and is sent to the deep tectal neuropil. Both spontaneous swims and visual stimulation alone are associated with comparable responses. However, when both coincide, the response magnitude is increased approximately linearly on the population level. Because the signal is mixed, apparent visual tuning can be confounded by motor events.

### Integration of visual and motor signals and synchrony in PrDA neurons

On average, the summation of visual and motor inputs was additive in driving PrDA responses at the population level (synergy ≈ 0, Fig. 5D). Interestingly, a similar mixing of input signals from distinct (sensory) channels has recently been reported for thalamocortical-like circuitry in juvenile zebrafish (Trinh et al. 2026). Notably, PrDA neurons exhibited highly synchronized population-level activity, even in the absence of any visual or motor drive (Fig. 6). This correlation was abolished by temporal shuffling, indicating that tonic coupling is an intrinsic property of this population. Such baseline synchrony may reflect shared recurrent excitation via local pretectal interneurons (Kubo et al. 2014, Zylbertal and Bianco 2023), commissural connections between the two hemi-clusters, and electrical synapses as described for dopaminergic neurons in the substantia nigra (Vandecasteele et al. 2005). On top of this intrinsic coupling, shared convergent visual and motor input could then drive the event-dependent increases in synchrony, so that baseline coupling sets a floor and common input scales synchrony above it.

The spatially uniform distribution of visual- and motor-related responses across the PrDA cluster (Fig. 3C, 4E) supports the idea that these neurons share a common input structure that naturally promotes co-activation. Visual and motor events progressively elevated this baseline synchrony, mirroring the single-neuron response pattern, with the strongest co-activation during coincident visual and motor events. This parallel scaling suggests that population synchrony and single-neuron gain share a common mechanism, that is, the summation of convergent inputs onto a recurrently coupled population. A similar event-dependent modulation has been described in primate dopaminergic systems, where noise correlations between midbrain DA neurons transiently increase following salient events and are thought to enhance the impact of dopaminergic signaling on downstream circuits (Joshua et al. 2009). Notably, synchronized activity has been observed also in other DA populations in larval zebrafish. Thus, pairs of DA neurons in the diencephalic posterior tuberculum exhibit phasic burst firing correlated with locomotor state (Jay et al. 2015), and many diencephalic dopaminergic clusters exhibit synchronized activity even during motor inactivity (Reinig et al. 2017). More broadly, the combination of high baseline coupling with context-dependent amplification of synchrony may be a general organizational principle of compact dopaminergic nuclei that broadcast modulatory signals to downstream circuits. Lastly, PrDA neuron activity may also be regulated by their expression of D2-like dopamine receptors (Fig. 1H). Because D2-like receptors signal through inhibitory G-proteins, this pattern is consistent with a negative feedback mechanism that limits neuronal excitability and transmitter release (Wolf and Roth 1990, Ford 2014). In PrDA neurons, such a mechanism may provide intrinsic stabilization of dopaminergic signaling during sustained or repeated behavioral activity.

More generally, the mixing of visual and motor signals in a predominantly visual region reflects a broader organizational principle, found across vertebrate brains. Many sensory areas have activity that is influenced by signals related to locomotion and behavioral states. For example, in mouse visual cortex, a high-dimensional component of the population activity tracks spontaneous movements (Stringer et al. 2019). Furthermore, the largest share of single-trial variance across the cortex is often attributed to movement rather than to stimulus or task (Musall et al. 2019). For the PrDA neurons investigated here, the mixed visuomotor signal reports a combination of visual and motor drive that may gate or bias the visuomotor transformation in the tectum. Between neurons, however, integration varied widely, spanning the range from sub- to supra-linear (Fig. 5D). A population with an additive mean response, but whose members range from suppressive to supra-linear boosting, can report the sum robustly while individual neurons may represent also different input mixtures. In this scenario, the between-neuron variation ranging from sub- to supra-linear summation may not be just heterogeneity in responses, but could in principle serve as a mixed representation of the sensorimotor state that can expand the range of variable combinations a downstream circuit can read out (Rigotti et al. 2013).

The origin of the visual and motor signals converging on PrDA neurons remains to be resolved. The close spatial apposition of PrDA dendrites with AF9 (Fig. 1D) suggests that PrDA neurons receive direct retinal input from a subset of RGCs (Burrill and Easter 1994). Such input could account for PrDA responses to both luminance increments and decrements, including bright and dark looms and flashes, as distinct subregions of AF9 exhibit ON-, OFF-, and ON-OFF-associated neural activity (Robles et al. 2014). However, monocular visual stimulation alone evoked bilateral activity (Fig. 4E), indicating retinal input cannot be the sole source for visual drive. This points to an additional commissural or interhemispheric contribution, as proposed for pretectal direction-selective neurons and for binocular integration in the tectum (Kubo et al. 2014, Gebhardt et al. 2019). Because PrDA neurons are active during fictive swims in immobilized larvae, their motor-related activity cannot arise from proprioceptive feedback, and instead likely reflects an efference copy from hindbrain premotor networks. However, resolving the presynaptic partners of PrDA neurons unequivocally will require connectomic reconstruction (Svara et al. 2018, Odstrcil et al. 2022, Svara et al. 2022).

### Implications of integrated PrDA signaling for tectal sensorimotor processing

Overall, the impact of stimulus- and motor-dependent DA release in the tectum likely depends on cell type-specific expression levels of DA receptors and their laminar organization. In the mammalian superior colliculus, D1-like receptors are enriched in superficial visual layers, whereas D2-like receptors are concentrated in deeper multimodal layers, producing layer-specific effects of DA (Bolton et al. 2015). Whether a comparable receptor topography exists in the zebrafish tectum is unresolved, but such an arrangement would in principle allow PrDA projections across distinct tectal strata to differentially modulate computations such as enhancing sensory salience, initiating action, or selecting between competing motor programs. Based on our finding that PrDA axons preferentially innervate deeper, non-visual layers, including the SGC and SAC, which are associated with tectal output pathways (Sato et al. 2007, Scott and Baier 2009), the dynamically regulated release of DA is likely to mainly modulate late steps of tectal visuomotor processing.

What, then, might be the functional impact of stimulus- and motor-dependent DA signaling in the tectum? First, by regulating excitability of tectal neurons during behavior, it may implement gain control along distinct tectal output pathways and dynamically adjust the threshold by which tectal representations are read out and transformed into motor commands. This view is consistent with work in lamprey showing that visually driven DA release enhances or decreases the excitability of tectal cells via D1- and D2-like receptors, respectively, thereby modulating the strength of orienting and escape responses (Perez-Fernandez et al. 2017). Second, because PrDA neurons exhibit phasic responses during self-generated motor output, their activity represents a corollary discharge signal (Crapse and Sommer 2008) that can inform the tectum about ongoing motor activity. Indeed, in earlier work we found that a phasic GABAergic corollary discharge signal effectively suppresses tectal projection neurons during saccade-like locomotion (Ali et al. 2023), mirroring the perceptual phenomenon of ‘saccadic suppression’ in mammals. In this context, the co-expression of *gad1* in PrDA neurons (Fig. 1H) may suggest that, through co-release of GABA, these neurons may be the source for motor-associated GABAergic inhibition during self-generated movement. However, whether PrDA neurons in fact release GABA and whether both DA and GABA are released onto the same neurons remain unresolved as dopaminergic neurons can restrict co-release of a second transmitter to only a subset of their targets (von Twickel et al. 2019). Furthermore, in the earlier work (Ali et al. 2023), GABA-mediated inhibition was associated with motor activity, but not with visual stimuli, arguing against PrDA neurons being the main source for motor-associated, phasic inhibition. Alternatively, assuming that PrDA action in the tectum is mainly mediated by DA, the integrated stimulus- and motor-dependent signal could represent a component of an internal model of the sensory consequences of the animal’s own movement. In this case, the convergence of sensory and motor information in PrDA neurons could result in a dopaminergic error signal that guides refinement of synaptic strength and connectivity in the developing retinotectal network and supports motor learning on slower timescales. In a similar vein, the integrated signal could mark those sensory representations that actually led to an action, since DA is known to regulate neuronal excitability and synaptic plasticity on behaviorally relevant timescales in reinforcement learning (Schultz 2007). Finally, as dopaminergic signaling in the tectum has been reported to shape tectal responses according to feeding state and metabolic demand (Zaupa et al. 2024), a further role of the PrDA pathway could be to regulate tectal sensory representations on longer time scales through DA-dependent changes in gene expression. Overall, these possibilities highlight the dopaminergic pretecto-tectal pathway as a potentially versatile modulatory influence on tectal sensorimotor processing, but resolving its precise functional role will require more research that directly addresses receptor-specific, transmitter-specific and cell-type specific mechanisms.

### Accounting for motor variables in visuomotor circuits

The sensitivity of PrDA neurons to both visual and motor-related events indicates that estimates of neuronal stimulus selectivity can be biased when differences in the efficacy of visual stimuli to drive motor activity are not considered. Thus, we observed that the apparent directional bias of PrDA neurons for forward moving gratings was abolished after accounting for swim-associated activity, suggesting that motor-associated signals can significantly affect estimates of apparent visual stimulus selectivity. This aspect could also affect estimates of neuronal direction selectivity in other pretectal populations and, similarly, extend to other visuomotor circuits, including those in the optic tectum itself. Together, this suggests that recordings of neural responses should ideally be paired with simultaneous measurement and analysis of motor variables in order to provide a more accurate basis for classifying neuronal response properties.

### Concluding remarks

In summary, our findings identify PrDA neurons as a sensorimotor dopaminergic pathway linking visual and motor circuits in the larval zebrafish diencephalon. These neurons integrate visual and locomotor-related signals, with enhanced activity during coincident sensory and motor events, positioning them to dynamically regulate tectal processing during behavior. More broadly, the convergence of sensory, motor, and modulatory signals onto tectal circuits may represent a conserved vertebrate principle for transforming visual information into adaptive behavior.

## METHODS

### Zebrafish husbandry

Zebrafish larvae (*Danio rerio*) were raised at 28.5°C in embryo medium in a 14h/10h light/dark cycle. Experiments were performed on zebrafish larvae 6-8 days post-fertilization, before sexual differentiation. The following transgenic lines were used: *Tg(th:th-e2A-QF2)m1512Tg*, *Tg(QUASr:GFP)c403Tg, Tg(QUAS:GCaMP6s)m1527Tg, Tg(atoh7:Gal4-VP16)s1992Tg and Tg(UAS:RFP)nkuasrfp1aTg* in the *nacre(mitfa-/-)* background. Animal husbandry and experimental procedures were performed following the guidelines of the German animal welfare law and approved by the local authorities (Regierungspräsidium Freiburg).

### *In situ* hybridization chain reaction (HCR)

All *in situ* HCR experiments were performed on 5 dpf *Tg(th:th-e2A-QF2)*; *Tg(QUASr:GFP)* transgenic larvae in the *mitfa*-/-background, treated with 0.003% of 1-phenyl2-thiourea (PTU) after collection and sorting of fertilized embryos. HCR reagents, namely, hybridization buffer, wash buffer and amplification buffer were purchased from Molecular Instruments. The ssDNA oligos were designed using a custom python script (Kuehn et al. 2022). Afterwards, the generated sequences were manually verified using A Plasmid Editor (APE), and at least 10 probe pairs evenly distributed along the gene were selected to be ordered from Sigma Aldrich. HCR probes for *th*, *dat*, *vmat2*, *gad1a/b*, *vglut2a/b*, *drd1a/b*, *drd2a/b* were used in this study. A probe set mixture was made by taking equal volumes of all the probes and mixed together.

The staining was performed according to a modified protocol of the manufacturer’s protocol (Molecular Instruments Inc.). Briefly, the larvae were anaesthetized in 1.5 mM tricaine and kept for overnight fixation in 4% paraformaldehyde solution in phosphate-buffer saline (PBS) at 4°C on a rotation wheel. Upon incubation, the larvae were washed 3x for 5 min with PBST (1x PBS + 0.1% Tween-20), followed by serial 10 minute treatments of 25% methanol/75% PBST, 50% methanol/50% PBST, 75% methanol/25% PBST, 100% methanol for dehydration. Afterwards, subsequent rehydration was performed in the reverse order of the dehydration steps and finally the larvae were washed for 5x 5 minutes in PBST. Ten larvae were transferred into a 1.5 ml tube and pre-hybridized with pre-warmed hybridization buffer for 30 minutes at 37°C. The probe solution was prepared by transferring 1 μl of 1 μM stock of probe set mixture to 500 μl of hybridization buffer at 37°C. The hybridization buffer was replaced with probe solution, and the samples were incubated for 12–16 h at 37°C with gentle shaking. Following incubation, the excess probes were removed by washing 4x 15 minutes with 500 μl of pre-warmed probe wash buffer at 37°C. Thereafter, larvae were washed 2× 5 min with 5× SSCT (5× sodium chloride sodium citrate + 0.1% Tween-20) buffer at room temperature in a flat shaker. Next, the larvae were incubated for 30 minutes at room temperature in 500 μl of amplification buffer for pre-amplification. Meanwhile, 30 pmol of hairpin h1 and 30 pmol of hairpin h2 (each from a 3 μM stock) were snap-cooled in separate PCR tubes at 95°C for 90 s in a thermocycler and cooled for 30 minutes in a dark environment. Afterwards, hairpin solution was prepared by transferring the h1 and h2 hairpins (each corresponding to the respective initiator sequence for the specific gene and tagged with Alexa Fluor 546 conjugate fluorescence protein) to 500 μl amplification buffer, which was added to the samples by replacing the pre-amplification buffer, and kept for 12-16 h in the dark at room temperature. Excess hairpins were washed the next day using 5× SSCT at room temperature as follows: 2x 5 minutes, 2x 30 minutes, 1x 5minutes. Larvae were then stored long-term at 4°C in 5× SSCT until imaging.

### Confocal imaging of transgenic larvae and HCR-labeled samples

Larvae were embedded in 1.6% low-melting agarose in 1x PBS. A laser scanning confocal microscope (FluoView1000, Olympus, with excitation wavelengths of 488 and 561 nm, respectively) with a water-immersion objective lens (Olympus 20x/1.0 NA) was used to acquire image stacks of PrDA neuron clusters, tectal hemispheres and retinal arborization fields.

### Image analysis for *in situ* HCR staining

Individual optical sections were taken at 5-µm steps from volumetric image stacks containing the PrDA cell cluster. Regions of interest (ROIs) corresponding to individual PrDA somata were detected from the GFP-channel using CellPose (Stringer et al. 2021) and subsequently manually proofread using ImageJ. Subsequently, the resulting ROI sets and corresponding optical sections from the HCR channel were analysed using Matlab (Mathworks). First, background fluorescence was estimated in individual optical sections of the HCR channel as median intensity of all pixels in that section and subtracted. Next, to identify transcript-positive subregions in a given optical section of the HCR channel, the HCR image was binarized using a Maximum Entropy method (Kapur et al. 1985): after normalization of the intensity histogram, the threshold T* for binarization was calculated using the formula:

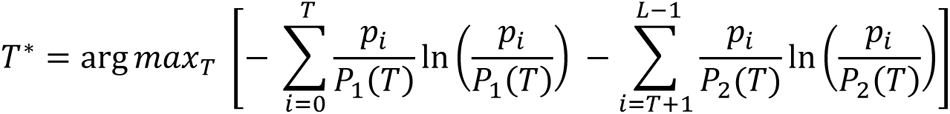

where p_i i_s the normalized intensity histogram, P_1 a_n P_2 a_re the cumulative probability distributions below and above a threshold T, respectively, and L the maximum intensity value in that image. Then T* is the intensity threshold value that maximizes the total entropy, defined as the sum of the background and foreground entropies. For calculation of T*, only pixels contained within the union of all soma ROIs was used in order to exclude spurious signals from non-cellular structures, such as skin. Next, soma ROIs were classified as transcript-positive if they contained one or more pixels with intensity values >T*. Finally, the proportion of transcript-positive somata was calculated as the number of transcript-positive soma ROIs divided by the total number of soma ROIs in the optical slice and reported as a percentage (Fig. 1H).

### Two-photon Ca2+ imaging

For Ca2+ imaging of neural activity in the PrDA neurons, zebrafish larvae were anesthetized using 0.02% MS-222 (tricaine methane-sulfonate, Sigma Aldrich) and then immobilized by incubation in α-bungarotoxin (1.0 mg/ml dissolved in embryo medium, Tocris). The larva was transferred to a custom-made cylindrical recording chamber (4 cm outer diameter, 2.5 cm height), with an extended arm to allocate and allow free movement of a borosilicate glass pipette for motor nerve recordings. The larvae were mounted in an upright position at approximately the center of the chamber on a sylgard-shelf that covered 10% of the bottom to leave an unobstructed view for visual stimulation. The chamber was filled with extracellular solution containing (in mM): NaCl (134), KCl (2.9), CaCl2 (2.1), MgCl2 (1.2), HEPES (10), and glucose (10), pH 7.8, Osmolality 290 mmol/kg.

Ca2+ imaging was performed using a multiphoton laser scanning microscope (Olympus FluoView FV1000), equipped with a water immersion objective (Olympus 20x/1.0 NA) and coupled with a Ti-Sapphire laser (MaiTai, SpectraPhysics) at an excitation wavelength of 920 nm. Emission light was filtered with a band-pass (515-560 nm). A rectangular region covering a single optical plane of PrDA neuron somata was imaged at ∼2.8 Hz throughout a 16 minute recording period during which different visual stimuli were applied (“sweep”). Consecutive sweeps were acquired in 5 µm steps, until the ventral edge of the PrDA cluster was reached.

### Visual stimulation

Visual stimuli were generated using VisionEgg (Straw 2008) and projected using a micro-projector (Kodak Luma 75) onto a diffusive screen (Rosco) covering the side wall of the recording chamber (Fig. 2A). Stimuli were presented monocularly to the left eye, spanning approx. 0° to 100° of the left visual field. Stimulus light was in the red-orange range, using a bandpass filter (606 ± 10 nm) positioned in front of the projector.

The stimulus protocol was structured as follows (compare Fig. 2): After 300 seconds with no stimulus light to record baseline spontaneous neuronal activity, a series of different stimulus types was applied: (1) Small white dot (5° visual angle) moving horizontally in a temporal–nasal–temporal direction at 20° elevation relative to the fish, repeated twice. (2) Abrupt appearance of a stationary vertical grating (spatial frequency: 20°, ‘grating appears’). (3) Grating moving temporonasally (‘grating forward’). (4) Grating moving nasotemporally (‘grating backward’). (5) Abrupt disappearance of grating (‘grating disappears’). (6) Expanding bright disc on dark background (exponential expansion, ‘bright loom’). (7) Expanding dark disk on a bright background (‘dark loom’). (8) Abrupt full-field illumination (‘bright flash’). (9) Abrupt full-field darkness (‘dark flash’). All stimuli were separated by 60 s inter-stimulus intervals to allow Ca2+ signals to return to near-baseline levels.

### Fictive swim recording

To record motor nerve activity from the tail corresonding to fictive swims, one suction electrode (open tip diameter ∼40 μm, borosilicate glass pipette with heat-polished tip; Science Products) was filled with extracellular solution and positioned on an intersegmental boundary in the midrange of the intact tail, and gentle suction was applied. Subsequently, motor nerve activity was recorded using an Axopatch 200B amplifier (Axon Instruments), and acquired at 10 kHz using LabVIEW (National Instruments, USA).

### Data preprocessing and analysis

Ca2+ imaging and motor nerve recordings were analyzed using custom scripts programmed in Python and R.

### Swim detection

Fictive swim bouts were detected using an automated procedure based on the envelope of the motor nerve signal (Suppl. Fig. 2A). First, the MNR signal was high-pass filtered (500 Hz) to remove slow drifts. Then we took the absolute value (rectification) and low-pass filtered it (zero-phase, 2^nd^-order Butterworth at 5 Hz), resulting in a zero-phase estimate of the smoothed amplitude. Next, the result was scaled by √2 for normalization and then squared to obtain the final envelope, representing signal power.

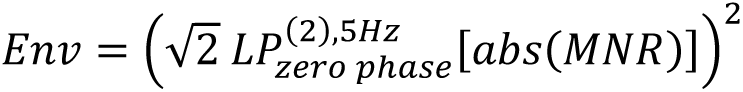

Swim bout onsets and offsets were detected by using a threshold of 1.5 times the standard deviation of the z-transformed envelope (Suppl. Fig. 2A). To increase detection accuracy, we defined additional criteria for detecting a swim bout. We checked the signal symmetry, quantified as the ratio between the absolute difference of positive and negative energy and total energy of the raw MNR signal.

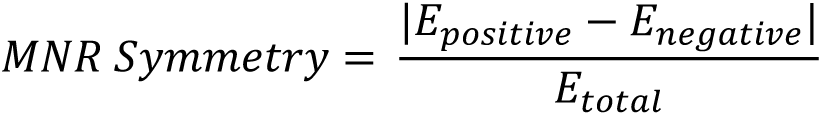

With the positive energy as the sum of all positive values squared and for the negative the same for all negative values (*E_pos_* = ∑*_x_*_>0_ *x*^2^ and *E_nenn_* = ∑*_x_*_<0_ *x*^2^).

Moreover, swim bouts shorter than 100 ms or longer than 2 s were excluded from analysis. Since the sampling rate and time dynamics of the Ca2+ recording cannot resolve the fast time scale of adjacent swim bouts, we merged bouts occurring within 8-10 s into single swim events. These swim events could then be related to neuronal activity on the time scales of Ca2+ imaging.

### Ca2+ imaging Analysis

Raw Ca2+ imaging data were corrected for motion artefacts using the non-rigid motion correction algorithm implemented in suite2p (Pachitariu et al. 2016). Based on standard deviation projections of the imaging time series, ROIs containing individual PrDA somata were selected, from which raw fluorescence signals were extracted using ImageJ. For each individual ROI, the raw signal was detrended and normalized using 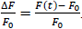. Baseline activity F_0 w_as computed using a running percentile window (10^th^ percentile) with a time window of 2 minutes. In total, signals from 360 neurons across 6 animals were analyzed.

### Response scoring

To characterize the responses of PrDA neurons, we utilized a regressor based scoring analysis approach (Miri et al. 2011, Förster et al. 2020). A regressor trace for all visual stimuli, representing an ideal Ca2+ response, was obtained by convolving a Ca2+ response function with a binary trace representing the stimulus onset times. The Ca2+ response kernel (CRK) was modeled as a double exponential function with a rise time constant τ_rise o_f 3 s and a decay time constant τ_decay o_f 6 s. These values were determined based on the Ca2+ imaging data by fitting a double exponential function to each detected Ca2+ transient and taking the median value of the resulting distributions (Suppl. Fig. 2C-F). For optimal fitting we included the amplitude A and a time alignment t_0._

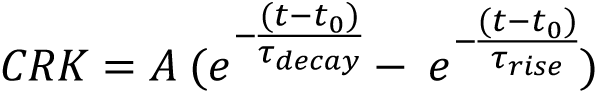

For scoring the ΔF/F response of a neuron to a stimulus presentation, a time window from 2 s before to 10-30 s after stimulus onset was used to fit an ordinary least squares linear regression between the stimulus regressor and the Ca2+ trace for each neuron and stimulus type (Suppl. Fig. 2I).

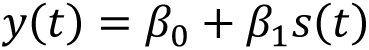

where *s(t)* is the specific stimulus regressor trace and *β_0_* _a_nd *β_1_* _t_he intercept and slope of the linear model, respectively.

Beforehand, the optimal time alignment was found using a cross-correlation between the two signals allowing for a stimulus-dependent time shift of 5 to 10 seconds (Suppl. Fig. 2G,H).

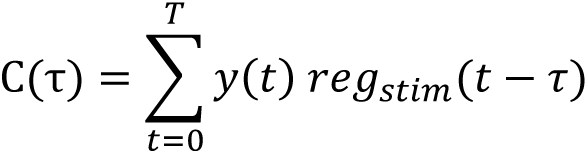

where *y(t)* being the measured ΔF/F trace and *reg_stim(_t)* being the respective regressor for a stimulus type. By multiplying the slope (*β_1_* _c_oefficient) and the *R²* value of the linear model, we assigned a score to each stimulus presentation (score = *β_1_* _×_ *R²*). This metric integrates response strength (slope *β_1_*_)_ with response reliability (goodness of fit, *R²*), providing a more robust measure of stimulus-related activity than either parameter alone.

The same procedure of creating regressor traces was applied to all detected swim events, yielding a motor regressor for each recording. For comparison between neurons from different animals, all scores of an animal were normalized by the 99^th^ percentile score of that animal (compare Fig. 5A).

To assess the significance of responses, we generated a null distribution for each neuron by computing scores at randomly chosen regressor times (10,000 iterations). The 95^th^ percentile of this null distribution served as the significance threshold (Suppl. Fig. 2J,K). Any event with a score above either the score threshold or the R^2^ threshold was considered significant.

For visualization purposes (Fig. 4B), we removed the putative contribution of motor events in the Ca2+ traces of all neurons by using the motor regressors to attenuate the Ca2+ signal. To attenuate motor-related components, each Ca2+ trace was divided by a scaled motor regressor and subsequently low-pass filtered at 0.5 Hz to remove sharp edges (Fig. 4B, bottom). The scaling factor of 100 was determined empirically, so that the attenuation effectively suppresses Ca2+ transients associated with spontaneous motor events in the first 5 minutes of a sweep.

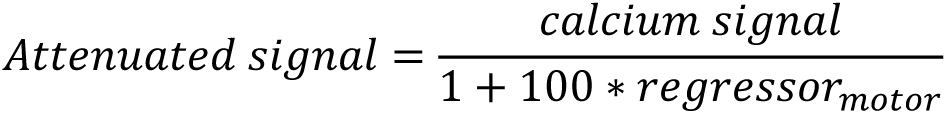

### Neuron classification

#### Stimulus specific classification

To classify the responses of PrDA neurons to different visual stimulus types we used a classification index (*CI _Stimulus_*, [−1,1]).

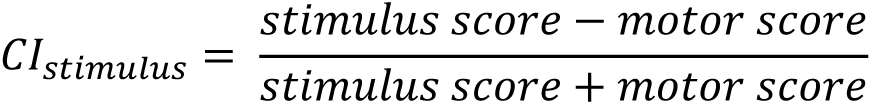

For each stimulus type, we computed such a *CI* for every neuron individually (Fig. 5B). The “stimulus score” is the result of the scoring between the stimulus regressor trace and the Ca2+ trace for that specific stimulus type and neuron. Respectively, the motor score is the result of the scoring between the motor regressor trace and the Ca2+ trace. Unlike the stimulus regressor, the shape of the motor regressor depends on an actual swim response happening during the stimulus phase. Thus, when there is no swim, then the motor score is zero. On the other hand, by design, there is always a non-zero stimulus regressor during visual stimulation, introducing an interdependency between the two scores. Therefore, a *CI_Stimulus_* _v_alue near zero indicates a mixed response, since the motor and the stimulus score are on a similar level. The neuron’s response to this stimulus type was, therefore, classified as “mixed”, since it is not possible to decide if the response is due to the stimulus or motor event. Following the same logic, a *CI_Stimulus_* _n_ear one indicates purely visual responses. Negative *CI_Stimulus_* _a_re conceptually not possible, since the motor score would need to be larger than the stimulus score, which is, due to their interdependency, not possible. By thresholding, we then classified the neuron’s response to a specific stimulus type as either “mixed”, “visual” or “no” response. The “no” response class was assigned when the stimulus score was not significant for a specific neuron and stimulus type. When the *CI_Stimulus_* _o_f a neuron was below 0.5, it was classified as “mixed”, otherwise as “visual”.

### Response type classification

The classification of neurons regarding their responsiveness to visual stimulation and motor events was evaluated by checking if a neuron had a significant response score for any purely visual stimulus (V+) and a significant score for spontaneous motor events (M+). Neurons recorded in sweeps with fewer than three spontaneous motor events were classified as motor unknown (M?). When a neuron had no significant visual or motor score, it was classified as V- and M-, respectively. Together this resulted in four different classes. Neurons with activity associated with visual and motor events (M+/V+), neurons with activity only to motor events (M+/V-) or vice versa (M-/V+) and neurons that showed activity in response to visual stimuli but it could not be determined whether they responded to motor events or not (M?/V+).

### Summation of visual and motor inputs

To quantify how visual, motor, and combined visual–motor inputs affect response scores of PrDA neurons, we computed signed selectivity indices for each neuron. Such a normalized difference is often referred to as a Michelson-like contrast index (similar to the *CI* above):

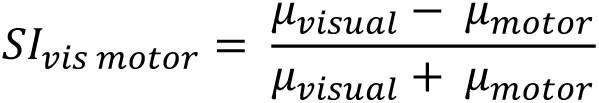

With *µ_visual_* _i_s the average of all purely visual stimulus scores (no motor events) for one neuron, and *µ_motor_* _i_s the average of all spontaneous motor event scores. All neurons recorded in sweeps with fewer than 3 spontaneous motor events were excluded in this analysis. Therefore, the *SI* compares the difference between the purely visual score and the purely motor score normalized by the sum of the two scores. If its value is above zero, the neuron’s response quantified by the scores for visual stimuli is larger than for spontaneous motor events. Below zero it is the opposite. Values around zero indicate equivalence. Similar to the *SI _vis motor_*, we computed also:

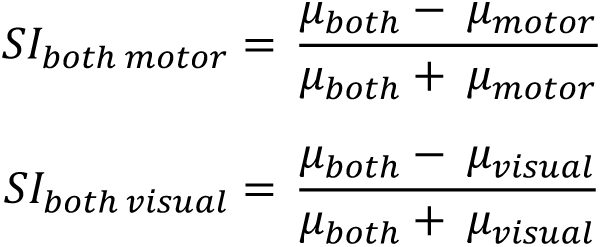

Here, ‘both’ means that both a visual stimulus and a motor event were present during the neurons response. Thus, *SI _both motor_* _a_nd *SI _both visual_* _a_sses if the neuron’s responses are larger when both inputs (visual stimulation and swim) are present together compared to either alone.

To specifically assess the input summation during simultaneous visual and motor events, we computed a signed synergy index:

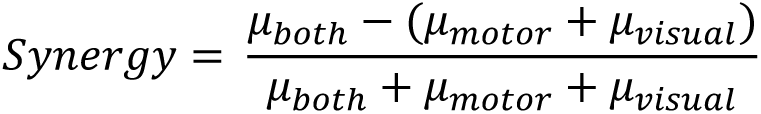

If motor and visual inputs are combined as a basic additive linear sum, the synergy index is around zero. A synergy index above zero indicates a supra-linear summation. A synergy index below zero indicates a sub-linear summation as in a signal suppression.

To avoid pseudo-replication and obtain non-parametric confidence intervals, we used hierarchical bootstrapping across fish to compute population-level summary statistics and significance tests. All selectivity and synergy indices were tested for being different to zero.

### Activity synchronization

To assess whether dopaminergic neurons in the pretectum exhibited synchronized activity, pairwise cross-correlations were computed between all pairs of neurons in each recording sweep. For every analysis window, each neuron’s activity trace was z-scored across time. For a lag *τ*, the cross-correlation between neurons (i) and (j) was calculated as

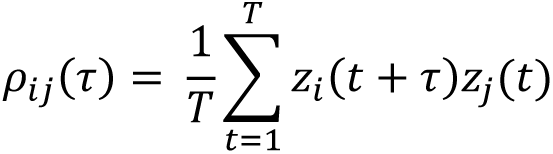

where *z_i(_t)* and *z_j(_t)* denote the z-scored activity traces and *T* is the number of overlapping time points at lag *τ*. Since the cross-correlation was computed on the z-scored activity, it approximates the Pearson’s correlation at each lag *τ*.

Activity traces were divided into 20 s windows corresponding to different conditions. These included spontaneous swim events, visual stimulus events with or without a swim response (both 10 s before to 10 s after onset) and spontaneous baseline windows of 20 s duration without nearby visual stimuli or swim events. For each condition, cross-correlations were computed for every neuron pair within each window and subsequently averaged across windows of the same condition (Fig. 6A, bottom panels).

To estimate the level of correlation expected by chance, shuffled control datasets were generated by independently circularly shifting each neuron’s activity trace by a random offset while preserving the temporal structure of the individual trace. Shift magnitudes were constrained to at least 20% of the window length to avoid trivial shifts. This procedure was repeated 200 times and plotted alongside the observed cross-correlations to provide a visual reference for the level of correlation expected by chance (Fig. 6A, right panels).

To quantify the recruitment on the population level, each sweep was analyzed independently. Within each recording, neuronal activity traces were first z-scored within each analysis window and then binarized by classifying a neuron as active whenever its z-scored activity exceeded 0.5. At each time point, the number of active neurons *k(t)* was determined. Recruitment was then defined as the fraction of active neurons at each time point,

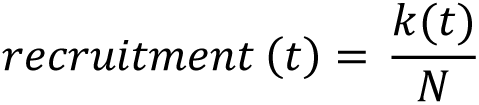

Where *N* denotes the total number of neurons in that recording. Thus, this metric ranges from 0 to 1 and represents the fraction of neurons that are active at the same time. Recruitment traces were computed for each sweep separately and subsequently aligned to event onset for averaging across events, recordings and larvae (Fig. 6B).

#### Direction selectivity

A behavioral direction selectivity index (*DSI_swim_*_)_ for drifting gratings was computed based on the number of swim bouts during presentation of a grating moving in the forward and backward direction, respectively (Fig. 7A).

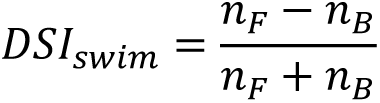

where *n_F_* _a_nd *n_B_* _a_re the number of swim bouts per stimulus presentation in the forward and backward direction, respectively.

An apparent neuronal direction selectivity index (*DSI_neuron,apparent_*_)_ for each neuron was computed correspondingly (Fig. 7B):

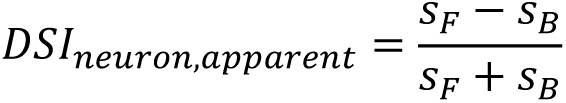

where *s_F_* _a_nd *s_B_* _a_re the stimulus regressor scores for forward grating and backward grating, respectively. Finally, in order to account for the apparent bias in the calculated stimulus scores due to differential motor-associated neural activity, we corrected each stimulus score with the number of swim bouts during visual stimulus motion to calculate a corrected neuronal direction selectivity index (Fig. 7C):

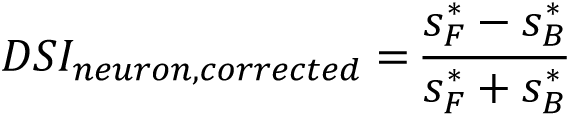

where

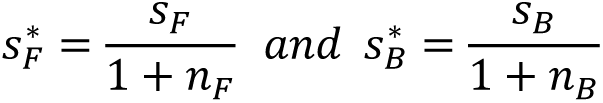

To test if the resulting distribution of direction selectivity indices differs from a uniform distribution we used a non-parametric Kolmogorov-Smirnov test.

#### Statistics

Statistical analyses were performed using Python, MATLAB, and R. Statistical tests are specified in the corresponding Method and Results sections and figure legends. Unless otherwise stated, statistical significance was set at p < 0.05.

## Acknowledgment

We thank Margit Böhler, Angela Wanninger and Sabine Götter for excellent support with fish husbandry and other technical help. Wolfgang Driever and Christian Altbürger for generously sharing transgenic lines. We thank Wolfgang Driever for comments on an earlier version of the manuscript and members of the Bollmann and Driever labs for helpful discussions. This work was performed with support from the German Research Association (DFG; Project-Nr. 446150760, 357057764, 453632629).

---

**Supplemental Figure 1:**
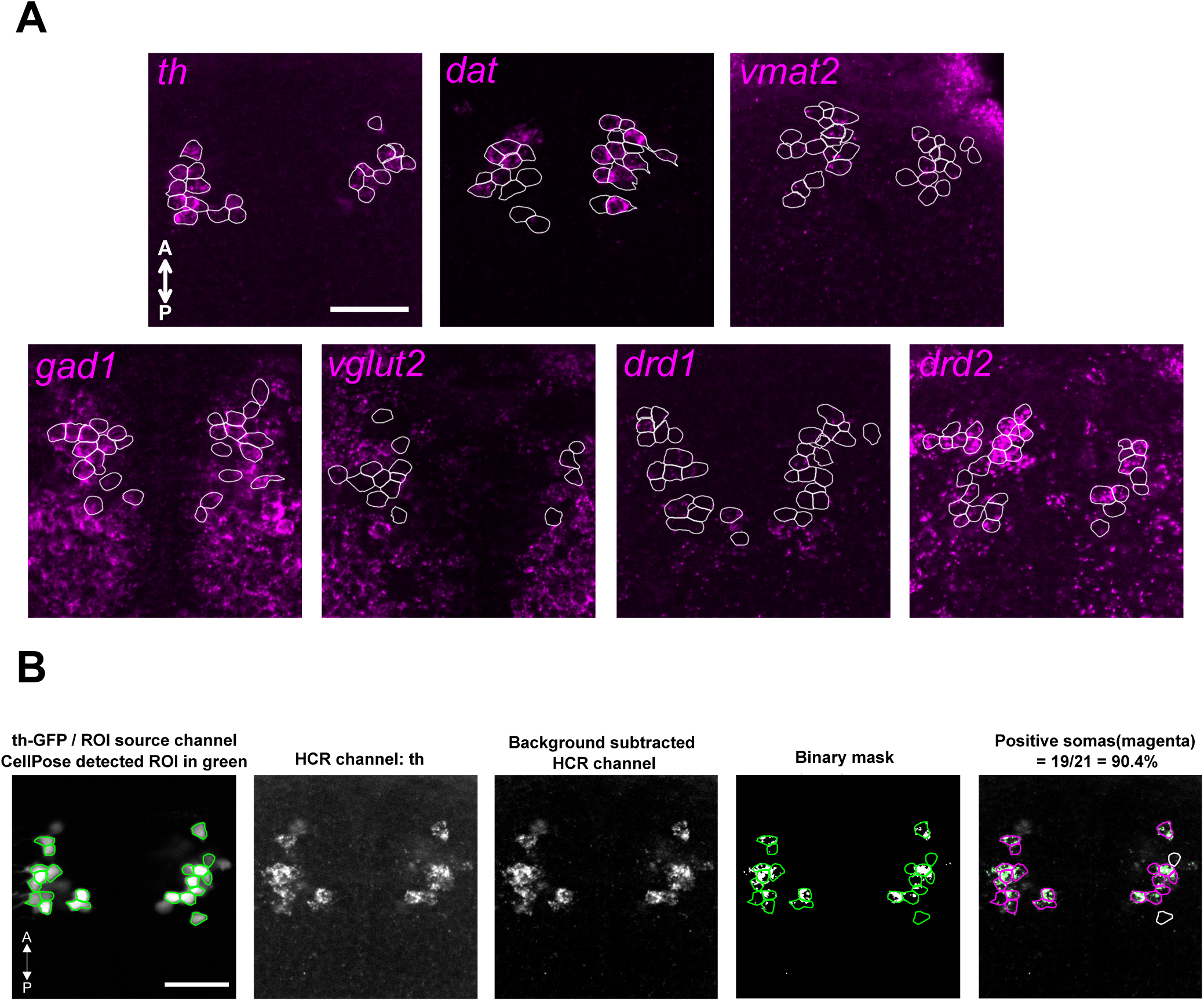
*In situ* HCR staining for marker genes and image processing workflow for quantification. A) Representative *in situ* HCR images showing staining of the dopaminergic markers *th*, *dat*, and *vmat2*, GABAergic marker *gad1*, glutamatergic marker *vglut2*, and the dopamine receptors *drd1* and *drd2* in pretectal dopaminergic neurons. White outlines indicate ROIs defined by GFP-positive PrDA somata. A, anterior; P, posterior. Scale bar 20 µm. B) Workflow for HCR transcript detection and quantification in PrDA neurons. *th*-expressing PrDA somata were identified from the GFP channel using Cellpose and outlined in green. The raw HCR channel, background-subtracted image, thresholded binary mask, and final classification of transcript-positive somata are shown. Magenta outlines indicate *th-*transcript positive somata, whereas white outlines indicate transcript-negative somata.

**Supplemental Figure 2:**
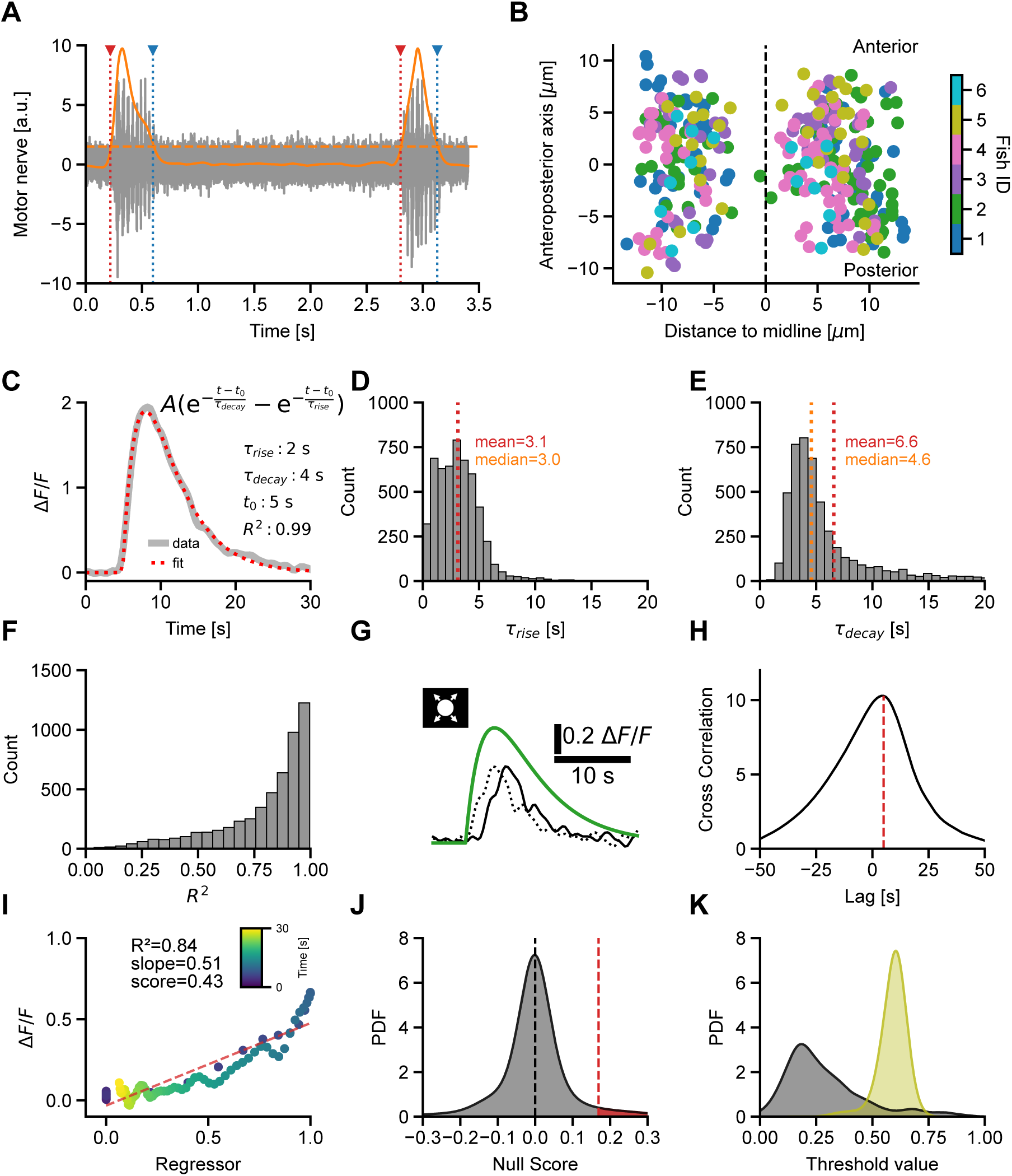
Functional imaging of PrDA neurons. A) Swim-bout detection algorithm. An envelope (orange) is computed from the MNR trace (gray, see Methods for details). Using a standard-deviation threshold, swim-bout onset (red) and offset (blue) are calculated from the envelope signal. B) Anatomical distribution of all recorded neurons from all larvae. Fish identity is color-coded. Black dashed line indicates the midline of the larva. C) Example Ca2+ transient (gray line), overlaid with a fit curve (red line) to determine time constants (τ_rise,_ τ_decay)_, for calculating a realistic Ca2+-response kernel (see equation) used in the scoring algorithm. D) Distribution of rise time constants τ_rise f_rom fits to all measured Ca2+ transients. Mean (red) and median (orange) values are indicated by vertical dotted lines. E) Distribution of decay time constants τ_decay f_rom fits to all measured Ca2+ transients. Mean (red) and median (orange) values are indicated by vertical dotted lines. F) Distribution of R² values for all exponential fits. G) ΔF/F transient (black) in response to a bright loom stimulus and the corresponding stimulus regressor (green). The dotted trace shows the ΔF/F transient after temporal alignment for optimal correlation (see H). H) Cross-correlation between the ΔF/F transient and the stimulus regressor as a function of time lag. The red dashed line indicates the lag (in seconds) yielding maximal correlation, used for temporal alignment in G. I) Scatter plot showing ΔF/F values versus regressor values over time (color-coded by time). The relationship is fitted using an ordinary least squares linear model, and neuron responsiveness is quantified as score = R² × slope. The red dashed line represents the fitted regression. J) Determination of significance thresholds for response scores. For each neuron, a null-score distribution was generated by randomly shuffling the onset times of regressors. Shown is one example neuron: the black dotted line indicates the mean value near zero, and response scores exceeding the 95th percentile of the null distribution (red area) were considered significant. K) Distribution of all threshold score values (gray) and corresponding threshold R² values across neurons.

**Supplemental Figure 3:**
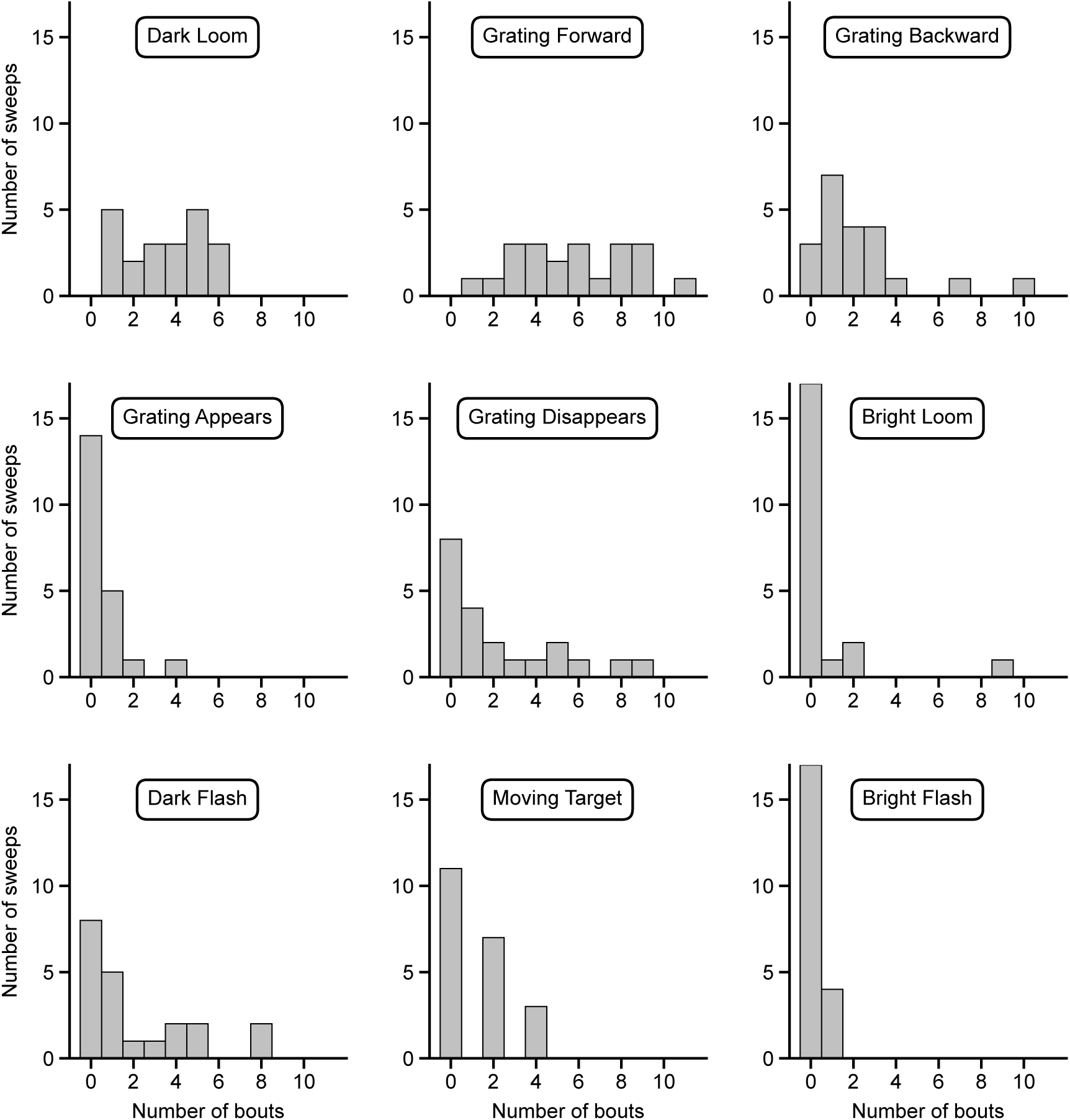
Number of triggered swim bouts per stimulus type. Histograms showing the number of swim bouts for each stimulus type.

